# Macrovascular contributions to resting-state fMRI signals: A comparison between EPI and bSSFP at 9.4 Tesla

**DOI:** 10.1101/2024.07.04.602062

**Authors:** Dana Ramadan, Sebastian Mueller, Ruediger Stirnberg, Dario Bosch, Philipp Ehses, Klaus Scheffler, Jonas Bause

## Abstract

The draining-vein bias of T2*-weighted sequences, like gradient echo echo-planar imaging (GRE-EPI), can limit the spatial specificity of functional MRI (fMRI). The underlying extravascular signal changes increase with field strength (B_0_) and the perpendicularity of draining veins to the main axis of B_0_, and are therefore particularly problematic at ultra-high field (UHF). In contrast, simulations showed that T2-weighted sequences are less affected by the draining-vein bias, depending on the amount of rephasing of extravascular signal. As large pial veins on the cortical surface follow the cortical folding tightly, their orientation can be approximated by the cortical orientation to 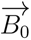. In our work, we compare the influence of the cortical orientation to 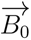 on the resting-state fMRI signal of three sequences aiming to understand their macrovascular contribution. While 2D GRE-EPI and 3D GRE-EPI (both T2*-weighted) showed a high dependence on the cortical orientation to 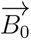, especially on the cortical surface, this was not the case for 3D balanced steady-state free precession (bSSFP) (T2/T1-weighted). Here, a slight increase of orientation dependence was shown in depths closest to WM. And while orientation dependence decreased with increased distance to the veins for both EPI sequences, no change in orientation dependence was observed in bSSFP. This indicates the low macrovascular contribution to the bSSFP signal, making it a promising sequence for layer fMRI at UHF.

## 1 Introduction

The carotid and vertebral arteries supply blood to the brain, branching into large arteries on the cortical surface, also called pial arteries. Diving intracortical arteries are oriented perpendicular to the cortical surface and large pial arteries. These diving vessels branch into randomly oriented arterioles (Duvernoy et al., 1981) and capillaries, where essentially gas exchange and nutrient delivery occurs for glia cells and neurons. The capillary branches connect to the venous blood stream through venules (∼10 *µ*m), which merge into intracortical ascending veins (∼80 *µ*m) oriented perpendicular to the large veins (≥ 200 *µ*m) on the cortical surface. The latter are referred to as pial or draining veins, as they drain the blood to the large cerebral veins of the brain. This means that both the arterial and venous vessels of the cortex follow the curvature closely. For this reason, the cortical orientation in respect to the main magnetic fields 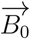 is a valid estimate for the pial vasculature. Like in previous work (Bàez-Yànez et al., 2017; Cohen-Adad et al., 2012; Gagnon et al., 2015; Viessmann et al., 2019), *θ_B_*_0_ is defined in this work as the angle between the cortical surface normal and the main magnetic field 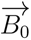.

It is known that the brain lacks an energy reservoir of its own, so blood flow is essential to supply the brain with the required energy on demand. When neurons are active, blood flow typically increases in their vicinity, and strong blood flow from activated larger areas accumulates in more distant larger veins. This phenomenon is referred to as neurovascular coupling (Attwell et al., 2010; Iadecola, 2004, 2017). For unknown reasons, active brain regions are flooded with more oxygenated blood than required (Raichle & Mintun, 2006). With this change in blood flow, blood volume and oxygenation level, also known as the hemodynamic response, the blood oxygenation level dependent (BOLD) effect arises (Ogawa et al., 1990), which is employed in the majority of functional magnetic resonance imaging (fMRI) experiments.

The deoxyhemoglobin in venous blood is paramagnetic and therefore perturbs the magnetic field in the surrounding diamagnetic tissue (Pauling & Coryell, 1936). If veins are modelled as infinitely long cylinders filled with paramagnetic material, the field offsets surrounding them are highest, when perpendicular to the main axis of 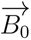 (Chu et al., 1990). Using this long cylinder simplification, the following equations can be used to describe the extravascular (Δ*ω_extra_*), as well as the intravascular (Δ*ω_intra_*) frequency offsets resulting from susceptibility differences between oxygenated and deoxygenated blood (Δ*χ*):

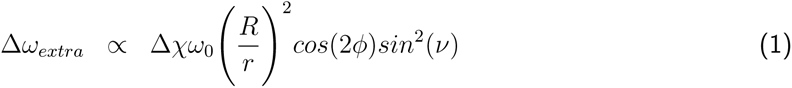

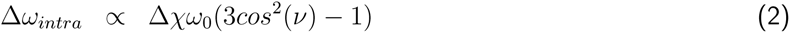

Here, *ν* is the angle between 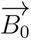 and the main axis of the cylinder, *R* its radius, *r* the distance between any observation point and the axis of the cylinder, and *ϕ* the polar angle of the position vector *r⃗* in the plane perpendicular to the cylinder axis. The Larmor frequency *ω*_0_ is directly proportional to B_0_, thus extra- and intravascular frequency offsets are stronger at higher field strengths. Since Δ*ω_extra_* in Equation 1 depends on the ratio 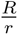, vessels with larger radius cause a more long-range field distortion than vessels with smaller radius. Due to larger spatial extent of this effect, the extravascular signal changes dominate at ultra-high field (UHF) strengths (Boxerman et al., 1995; Uğurbil et al., 1999; Uludağ et al., 2009).

During neuronal activation, excessive oxygenated blood flows to the activated area and through the venous system. The differences between the oxygenated and deoxygenated blood in the veins changes T_2_* times. This change is usually captured with gradient echo (GRE) sequences. In addition, higher blood flow and blood volume in an active brain area increases T_2_ times due to higher diffusion of spins in this area. Diffusion effects are more prominent around small vessels, because the diffusion distance relative to the vessel size is higher (Weisskoff et al., 1994). Spin echo (SE) sequences highlight this effect. They are not sensitive to spin dephasing around veins with a diameter way larger than the diffusion length, due to their refocusing pulse. With increasing field strength, the T_2_ effect is increased relative to the T_2_* contrast (Yacoub et al., 2001). Thus, it can be concluded that SE sequences are more sensitive to smaller veins and therefore more spatially specific, particularly at high and UHF strengths.

Echo-planar imaging (EPI) is the earliest sequence used for high-speed image acquisition (Mansfield, 1977; Stehling et al., 1993). It also has a high BOLD sensitivity, making it the most widely used readout in fMRI (Belliveau et al., 1991; Ogawa et al., 1990). However, it suffers from geometric distortion and susceptibility artifacts increasing with field strength due to B_0_ inhomogeneities (Moeller et al., 2010; Speck et al., 2008). Moreover, increased blurring effects are expected with increasing field strength due to faster T_2_* decay. The T_2_* weighting of GRE-EPI not only leads to a bias towards large draining veins on the cortical surface (Bause et al., 2020; Lai et al., 1993; Turner, 2002), and a loss of specificity towards the cortical surface in layer fMRI, but also to an increase in sensitivity (Polimeni et al., 2010). SE-EPI, on the other hand, is severely limited by the higher specific absorption rate (SAR), which can limit the temporal resolution and / or spatial coverage especially at field strengths ≥7 T (Budde et al., 2014). Therefore, GRE-EPI is still preferred at UHF strengths.

Fortunately, other sequences, like the balanced steady-state free precession (bSSFP), may also be utilized to map the BOLD effect at high and UHF strengths (Bowen et al., 2005; Miller, 2012; Scheffler & Ehses, 2016; K. Zhong et al., 2006). Although signal acquisition is slower compared to the EPI technique, resulting images do not suffer from geometric distortions. The bSSFP sequence is efficient as it maintains the magnetization by acquiring the signal in the steady state. Moreover, the contrast in bSSFP is T2/T1-weighted and although it is a GRE sequence, echo formation at TE = TR/2 (which is used in this work) is similar to SE, meaning that the transverse magnetization is (nearly) refocused when the echo is acquired (Scheffler & Hennig, 2003). In theory, the SE-like signal of bSSFP shows increased selectivity to the microvasculature at high fields (Bowen et al., 2005) with a high intravascular contribution (Pérez-Rodas et al., 2021). This theoretical expectation was further strengthened by Monte Carlo simulations using sections of vascular networks of the mouse parietal cortex (Bàez-Yànez et al., 2017). It can therefore be expected that bSSFP enables the acquisition of more specific fMRI signals, closer to the neuronal activation sites.

With increasing field strength (*B*_0_), the overall signal-to-noise ratio increases (Pohmann et al., 2016). Smaller voxel sizes are feasible, increasing the heterogeneity of voxels in tissue, vessels or CSF and reducing physiological signal confounds (Triantafyllou et al., 2005). Moreover, simulations have shown that signal contribution from the microvasculature increases by the square of *B*_0_ (Ogawa et al., 1993), while signal from the macrovasculature increases only linearly. Accordingly, a more specific BOLD effect should occur at UHF compared to lower field strengths. However, as the extravascular effect dominates at UHF, a higher signal contribution from large draining veins can be expected in T_2_*-sensitive sequences and thus an increased dependence of signal fluctuation on the cortical orientation, *θ_B_*, relative to 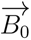. For example, Gagnon *et al*. showed that the distinct orientation of the cortical macrovasculature is not reflected in the SE signal amplitude (Gagnon et al., 2015) but in the GRE signal by performing simulations of the extravascular signals in vascular models and GRE BOLD measurements during hypercapnia. The concept was then further strengthened by Bàez Yàñez *et al*. by comparing the cortical and therefore vascular orientation dependence of GRE, SE and bSSFP in simulations. This study showed a BOLD signal change dependence on the cortical orientation that varied with cortical depths for all three sequences and a high signal sensitivity of bSSFP to the microvasculature, similar to SE (Bàez-Yànez et al., 2017). The strong orientation dependence of the GRE-EPI at UHF was then demonstrated by Viessmann *et al*. (Viessmann et al., 2019), who investigated the resting-state fMRI signal of 2D simultaneous multi-slice (SMS) GRE-EPI (Setsompop et al., 2012) at 3 T and 7 T. They observed a much stronger *θ_B_*_0_ -dependence of the fMRI signal fluctuation at 7 T compared to 3 T, calculated as the coefficient of temporal signal variation (CV). The angular dependence of CV followed a cos^2^(*θ_B_*) curve which can be expected for GRE-EPI from Equation 1, as 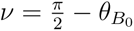 with *θ_B_* being the angle between the cortical surface normal and 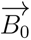.

For sequences with reduced macrovascular contribution, like bSSFP, the orientation dependence should be lower, since the intravascular effect dominates at 3 T and 9.4 T, especially in the macrovasculature and for short repetition times (TRs) (Pérez-Rodas et al., 2021).

To further investigate this phenomenon experimentally, we compare the resting-state fMRI signals of 3D bSSFP (Ehses & Scheffler, 2018) with segmented 3D GRE-EPI (Stirnberg & Stöcker, 2021) and 2D SMS GRE-EPI (vendor provided) in their cortical orientation and cortical depth dependence at 9.4 T. In the 2D SMS GRE-EPI examinations a similar sequence protocol like in a previous study (Viessmann et al., 2019) was applied to allow for better comparison of results at an even higher field strength. Furthermore, we investigated the dependence of the resting-state signal on the cortical orientation in relation to the distance to large veins, expecting a lower orientation dependence when investigating voxels further away from veins.

## 2 Methods

### 2.1 Data Acquisition

The presented study is part of a test plan for the development of 9.4 Tesla functional imaging methods reviewed and approved by the Ethics Committee of the University of Tübingen (approval no. 649/2021BO1). The volunteers provided their written informed consent to participate in this study.

MRI data of the brains of six subjects were acquired on a whole-body 9.4 Tesla scanner with an in-house built coil (16 Tx/ 31 Rx) (Shajan et al., 2014). Data from one subject were discarded due to excessive motion. For each subject, two sessions of resting-state fMRI measurements were acquired at the same time of day. In the first session, four runs of each 3D bSSFP (Ehses & Scheffler, 2018) and segmented 3D GRE-EPI (Stirnberg & Stöcker, 2021), both with a volume repetition time (TR_vol_) of 3 s, and one run of 2D SMS GRE-EPI with a TR_vol_ of 1.7 s were acquired. The imaging parameters of both 3D sequences were optimized to obtain comparable spatial coverage and TR_vol_. The acquisition time of about 5.5 min was held constant for all sequences by adjusting the number of measured frames. All functional images were acquired with an isotropic voxel resolution of 1.1 mm. The imaging slab of all sequences was oriented parallel to the calcarine sulcus, but almost whole brain coverage was only achieved with the 2D SMS protocol. The orientation and positioning of the slab was chosen to cover as many cortical orientations as possible. The resulting imaging coverage is shown in Figure 1. In the second session, four runs of 2D SMS GRE-EPI and one run of each 3D bSSFP and segmented 3D GRE-EPI were measured to test for test-retest variability. In both sessions, additional frames of both EPI sequences were acquired with the same parameters and reversed phase encoding (PE) direction to perform unwarping and co-registration of the anatomical data (explained in more detail in subsubsection 2.2.3). Acquisition parameters for all sequences are stated at the end of this section.

**Figure 1:**
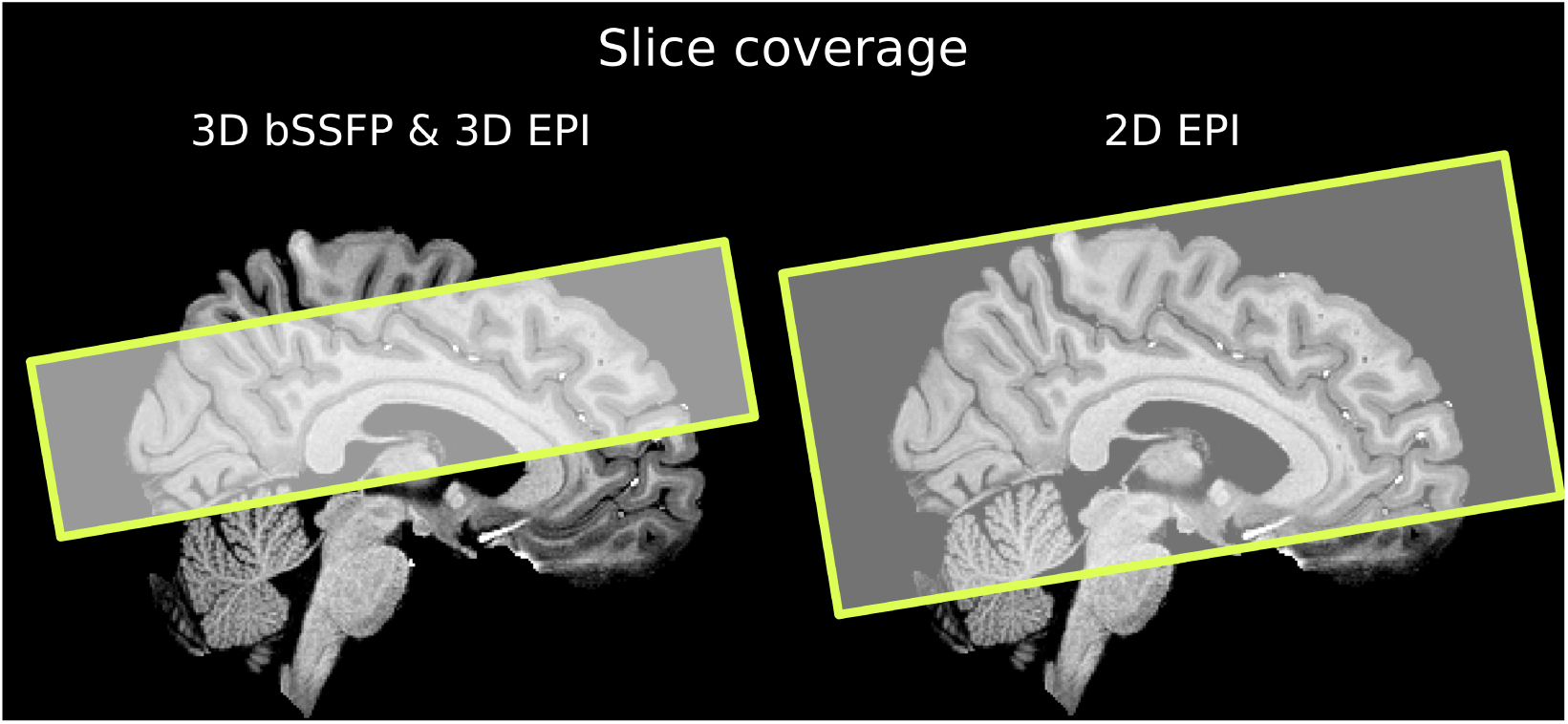
Slice coverage for the 3D and 2D sequences. A slab was acquired parallel to the calcarine sulcus, covering 40 and 44 slices for 3D bSSFP and 3D EPI, respectively. The same orientation and higher slice coverage (87 slices) was chosen for 2D EPI. The slice coverage is shown on a skull stripped, T1-weighted MPRAGE after bias field correction for subject S5. Similar slice orientation and coverage was chosen for the other subjects.

Physiological parameters were acquired with a respiration belt and photoplethysmograph for breathing and heart rate tracking, respectively (Biopac Systems Inc., CA, USA). However, physiological noise regression was not performed for all runs due to unstable recordings of the scanner’s trigger signal which hampered the temporal synchronization with the data. Runs with complete physiological recordings were evaluated.

CSF nulled whole-brain T1-weighted MPRAGE was acquired during only one session, usually at the end of the first one. For this sequence, a non-selective, population-optimized parallel transmit pulse was used (Bosch & Scheffler, 2023; Gras et al., 2017) in order to improve signal homogeneity. In a third session, 0.48 mm multi-echo (ME) GRE were acquired for the calculation of susceptibility-weighted images (SWI) (Haacke et al., 2010) to estimate the location of large veins. The sequence was modified to allow for the recording of an FID after the acquisition of the last spatially encoded echo for the retrospective correction of physiologically induced phase fluctuations (Vaculčiakovà et al., 2022). Unfortunately, one subject (S1) did not meet the requirements to be measured in the third session, which made data acquisition and corresponding analysis for S1 impossible.

The **acquisition parameters** were as follows:

- **3D bSSFP:** nominal matrix size: 174×174×40, number of volumes: 112, resolution: 1.1 mm isotropic, TR/TE = 3.14 ms/1.57 ms, TR_vol_ = 3000 ms, nominal flip angle (FA) = 11*^◦^*, slice partial Fourier (pF) = 6/8, GRAPPA recon (Griswold et al., 2002), R = 5×1, 10 % slice oversampling, readout (RO) bandwidth = 1250 Hz*/*px.
- **segmented 3D GRE-EPI (3D EPI):** nominal matrix size: 174×174×44, number of volumes: 112, resolution: 1.1 mm isotropic, TR_vol_ = 3000 ms, TE = 22 ms, FA = 15*^◦^*, slice pF = 6/8, GRAPPA recon, R = 3×1, PE_1_ = anterior-posterior (AP), PE_2_ = HF, interleaved multi-shot segmentation factor = 2, nominal echo spacing = 1.07 ms, RO bandwidth = 1106 Hz*/*px.
- **2D SMS GRE-EPI (2D EPI):** nominal matrix size: 174×174×87, number of volumes: 197, resolution: 1.1 mm isotropic, TR_vol_ = 1700 ms, TE = 25 ms, FA = 60*^◦^*, acceleration mode = slice acceleration, acceleration factor PE = 4, SMS = 3, PE direction = AP, nominal echo-spacing = 0.79 ms, RO bandwidth = 1512 Hz*/*px.
- **MPRAGE:** nominal matrix size: 256x306x300, resolution: 0.7 mm isotropic, TR_inversion_ = 3400 ms, TR_readout_ = 6.5 ms, slice pF = 6/8, GRAPPA recon, R= 2×1, FA = 7.5*^◦^*, TI = 1460 ms, bandwidth = 400 Hz*/*px.
- **ME-GRE for SWI:** nominal matrix size: 422×410×176, resolution: 0.48 mm isotropic, TR = 34 ms, TE_1_ = 7.58 ms, TE_2_ = 15.16 ms, TE_3_ = 22.74 ms, TE_4_ = 18.2 ms, FA = 16*^◦^*, GRAPPA recon, R = 2×2, RO bandwidth = 213 Hz*/*px.

### 2.2 Data Analysis

#### 2.2.1 Functional Data Processing

The first two volumes of the 3D EPI and 3D bSSFP measurements were removed to ensure that the analyzed data were acquired in the steady state. As the TR_vol_ of 1700 ms for the 2D EPI was shorter compared to the 3D sequences (3000 ms), three instead of two volumes were discarded. This resulted in 110 and 194 volumes for further processing of the 3D sequences and the 2D EPI, respectively.

Motion correction was performed in SPM12 (v7771) registering all volumes of all runs of one session to the mean volume of that session to minimize interpolation bias. Second degree B-spline interpolation and no up-sampling was used to perform this step. The full width at half maximum (FWHM) of the Gaussian smoothing kernel applied to the data to estimate realignment parameters was set to 1 mm.

The CV was calculated voxel-wise from the motion corrected volumes as a measure of TR-normalized time series signal fluctuation:

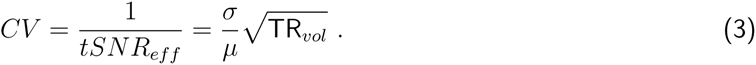

Here, tSNR_eff_ is the tSNR efficiency, given by the time series mean, *µ*, divided by the time series standard deviation, *σ*, normalized by the square root of the volume TR. This normalization was performed in order to better compare CV of the 3D sequences (TR_vol_ = 3 s) to the 2D EPI (TR_vol_ = 1.7 s).

Distortion correction was performed only on images acquired with the EPI sequences using FSL’s TopUp (6.0.5.2) (Andersson et al., 2003; Smith et al., 2004). The inversion of the resulting field map was used to warp the anatomical data and the values calculated in the anatomical space, namely (a) the orientation values in voxel space, (b) the ribbon file (gray matter (GM) - white matter (WM) segmentation), and (c) the vein distance maps to the respective EPI space (explained in more detail in subsubsection 2.2.3). It is important to note, that TopUp was only used to unwarp the EPI image temporarily and thus be able to co-register the anatomical image to the distortion corrected EPI and to obtain the field-maps to finally warp the anatomical images to the distorted functional space. All further analyses were performed in the functional space.

#### 2.2.2 Anatomical Data Processing

The T1-weighted MPRAGE was first bias field corrected in SPM12 using the segment tool, 40 mm FWHM and medium regularization. The bias field corrected images were then skull stripped to get a brainmask using the integrated synthstrip method in FreeSurfer mri synthstrip (Hoopes et al., 2022), as this skull-stripping method resulted in a better GM-WM segmentation compared to the regular one used in the recon-all pipeline of FreeSurfer (v.7.4.0). The latter pipeline was then used to get a segmented brain and mainly two surfaces, the white surface (between WM and GM) and the pial surface (between GM and the cerebrospinal fluid (CSF)). As surfaces consist of vertices, and three vertices form a closed surface, the surface normal from each small surface was calculated as explained in (Viessmann et al., 2019). The angle between the surface normal and 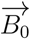 is defined here as *θ_B_*. These surface values were then transformed into voxel space at the native resolution of the MPRAGE of 0.7 mm.

#### 2.2.3 Co-registration and Depth Segmentation

The T1-weighted volume along with the cortical ribbon and orientation values calculated in FreeSurfer were co-registered to the functional volumes as follows: First, the T1-weighted MPRAGE was linearly co-registered to either the mean bSSFP volume or the distortion corrected mean EPI volume in SPM12 by maximizing the entropy correlation coefficient. Second, the cortical ribbon and the orientation values were transformed accordingly using the resulting transformation matrix. Third, an additional warping step using the calculated field map from TopUp was performed on all such co-registered anatomical data to match the distortions in the respective EPI data. The last step was skipped when matching the anatomical data to the bSSFP data, as the line-per-line acquisition does not result in susceptibility-induced distortions along the phase encode direction. After co-registration (and warping), the data were up-sampled by a factor of four in all three dimensions using nearest neighbor interpolation. Finally, five equi-distant cortical depths were calculated in LayNii (v.2.6.0) (Huber et al., 2021) from the co-registered (and warped) cortical ribbon.

#### 2.2.4 SWI and Vein Mask

The ME-GRE data were reconstructed offline using in-house developed Matlab (The MathWorks Inc., Natick, MA, USA) routines. The temporal variations of the phase measured by the FID were first subtracted from the corresponding k-space lines using the median FID phase over time as a reference. After GRAPPA reconstruction, complex image data were obtained using the adaptive combine approach (Walsh et al., 2000). Afterwards, brain extraction was performed from the resulting images using FreeSurfer mri synthstrip before SWIs were calculated by repetitive application of a sigmoid filter of the phase data (Santiesteban et al., 2006). The SWIs of the different echoes were then combined as a sum-of-squares to improve contrast, especially for small veins. The resulting, ’vessel-weighted’ image was used as an input for a Matlab implementation of the Jerman Vessel filter (Jerman et al., 2016). The resulting data were then thresholded before, in a final step, the vessel mask underwent careful manual inspection in FreeView using the second echo of the ME-GRE as a reference. Particularly at the cortical surface and between the cerebral hemispheres, it was necessary to remove falsely detected vessels. Note that by combining the different echo times, the resulting vascular mask was very conservative since veins appear larger in later echoes due to increased intravoxel dephasing.

Similarly as for the T1-weighted data, the vein mask and the ME-GRE were co-registered to the functional data: First, the second echo was co-registered to either the distortion corrected EPI volumes or the mean bSSFP image. The obtained transformation matrix was then used to re-sample the vein distance map to the (distortion-corrected) functional space. Lastly, these vein distance maps were warped in order to match the respective EPI-distortions in PE direction. Since the vein distance maps were provided in a continuous scale, the data were binned to include all voxels within high (0 mm to 0.71 mm), medium (0.71 mm to 1.41 mm) and low (1.41 mm to 2.12 mm) proximity to the veins. Closest voxels with a value of 0 mm are located inside the veins.

## 3 Results

### 3.1 Data Quality and Segmentation

Data quality is shown in Figure 2 for each sequence. An exemplary axial slice of one run from subject S5 and corresponding CV values are presented. It can be seen that 3D EPI show lowest CV values (highest tSNR efficiency), followed by 3D bSSFP and 2D EPI. For bSSFP, CV values in GM are lower than in WM, reflecting higher tSNR values in GM. All three sequences have CV values in similar ranges, which makes a comparison between the different sequences feasible.

**Figure 2:**
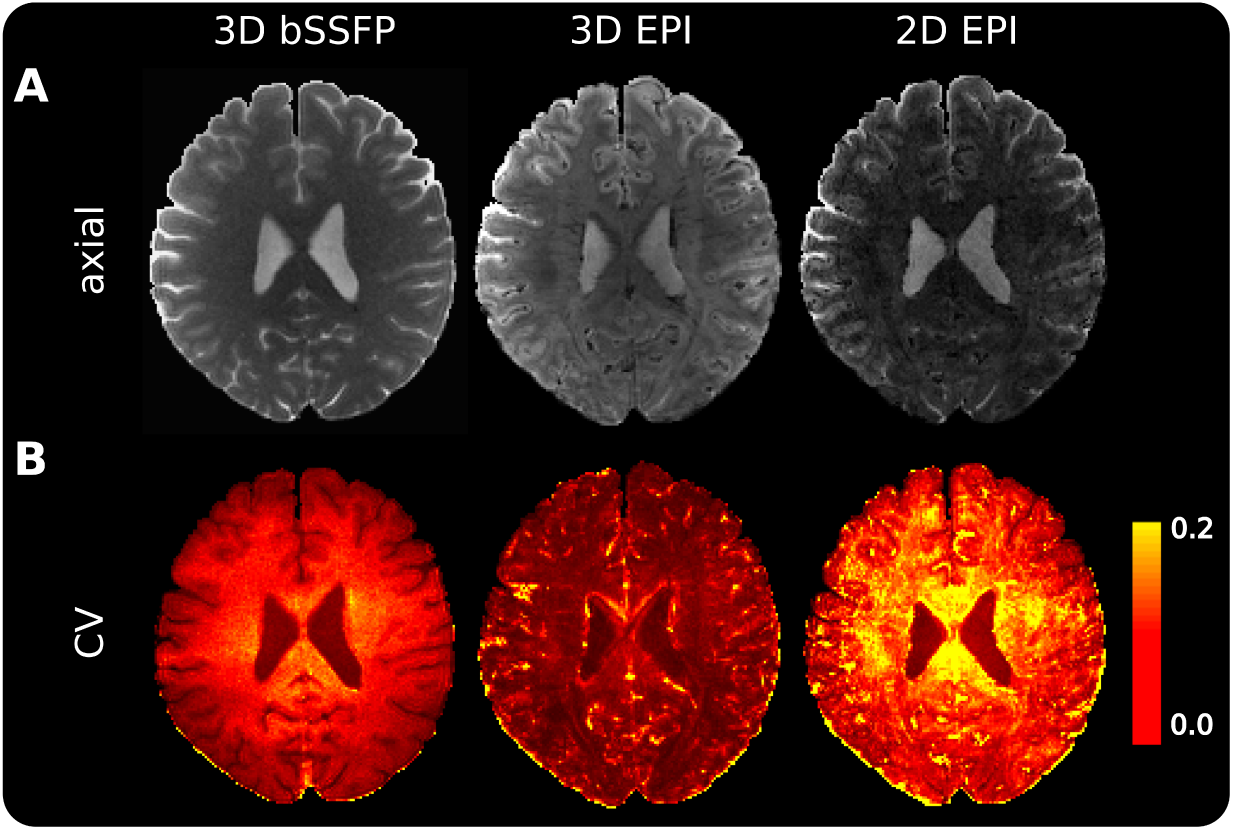
Exemplary axial slice of one volume to demonstrate data quality. (A) from one run of subject S5 for 3D bSSFP (left), segmented 3D EPI (middle) and 2D SMS EPI (right) before motion correction. (B) The CV values calculated from the time series of the respective sequences after motion correction.

Up-sampling is recommended before performing depth segmentation using LayNii (Huber et al., 2021) to obtain smoother equi-distant results. Moreover, LayNii works best with straight patches and performs poorer in curved regions. Therefore, we up-sampled the co-registered cortical ribbon by a factor of four to get voxel dimensions of 0.275 mm isotropic and be able to divide the ribbon into five equi-distant depths with Depth 1 (D1) and Depth 5 (D5) closest to CSF and WM, respectively. The difference between the original data and the up-sampled data using nearest neighbor interpolation can be seen in Figure 3 that shows the number of voxels per cortical depth without (A) and with (B) up-sampling. Note that every third slice in each dimension after up-sampling is considered, as otherwise data points would be repetitive. An uneven distribution of voxels (with more voxels in the middle layers) can be seen for the original data without up-sampling, both from the bar plots and from the overlaid depth profiles displayed below. Therefore, unless otherwise stated, all results shown are obtained after up-sampling.

**Figure 3:**
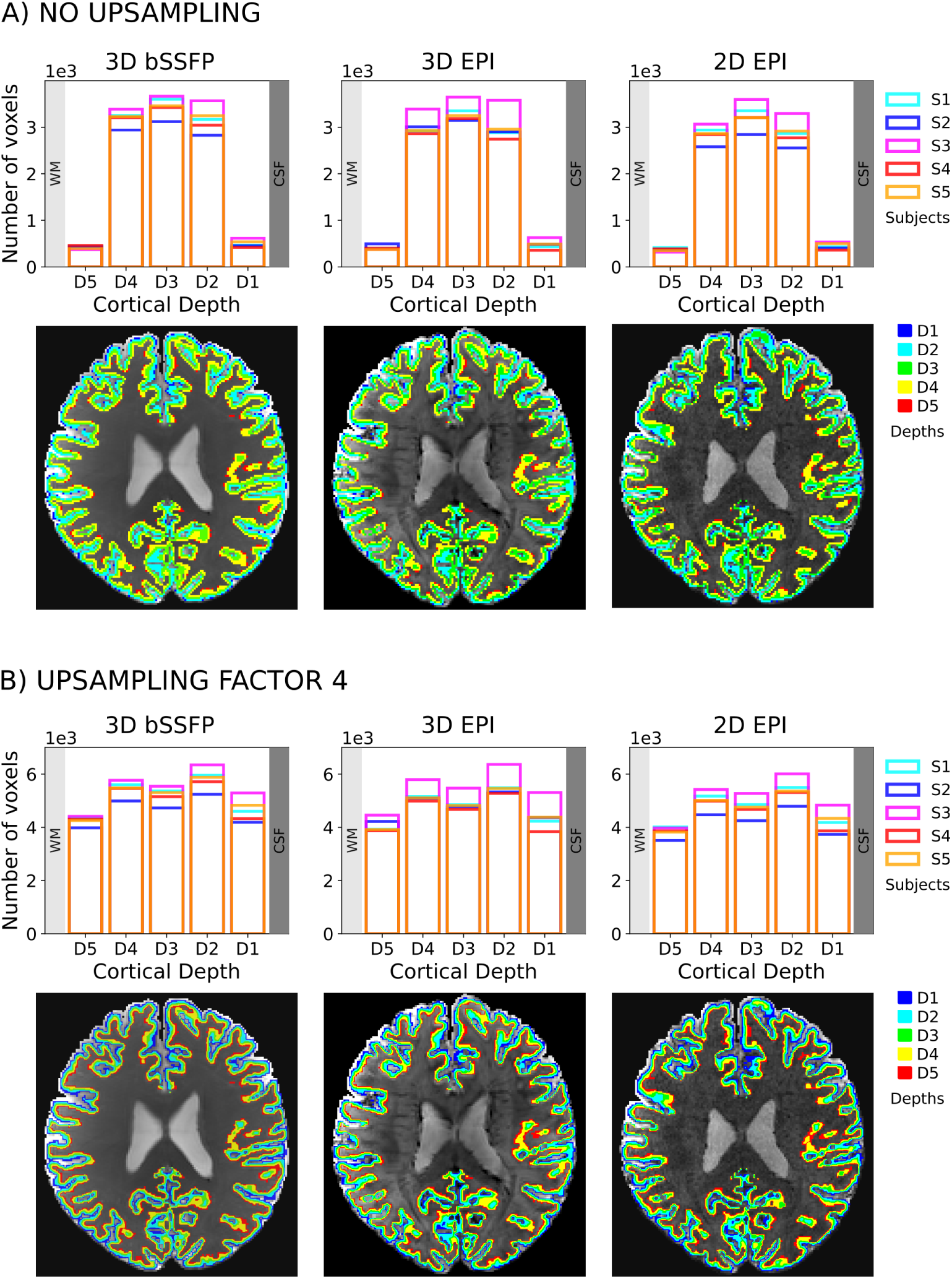
Distribution of voxels in the cortical depths without. (A) and with (B) up-sampling by a factor of four. Bar plots of the voxel counts are shown for all subjects (upper row) and the depths calculated with LayNii are overlaid on the mean functional image of one subject (lower row) for all sequences. The depths are more equally distributed after up-sampling.

The distribution of the cortical orientation values to 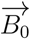 after co-registration (and warping) is shown in Figure 4A. It can be seen directly, that more voxels have a *θ_B_*_0_ value around 90°, i.e. are located in a cortical region with an orientation parallel to the main magnetic field. This is expected and can be explained by the distribution of angles for randomly located points on the surface of a sphere, which can be described by a sine curve. However, due to the unequal number of samples for the different angles, this distribution would introduce a strong bias into the analysis of the data. To remove this bias, the cortical orientation bin sizes are adjusted accordingly, so that each point in each plot of the following figures corresponds to a similar number of voxels. An example of how bins are adjusted can be seen in Figure 4B. Figure 4C shows the *θ_B_*_0_ values on the corresponding co-registered images. In the axial slices only the co-registration quality can be assessed. For that reason, we additionally show one coronal slice for 2D EPI with a zoomed-in version. It can clearly be seen, that the cortical orientation values correspond to the expected *θ_B_* values (blue on the medial gyrus (*θ_B_* = 0*^◦^*) and red in the more lateral sulcus (*θ_B_* = 90*^◦^*)).

**Figure 4:**
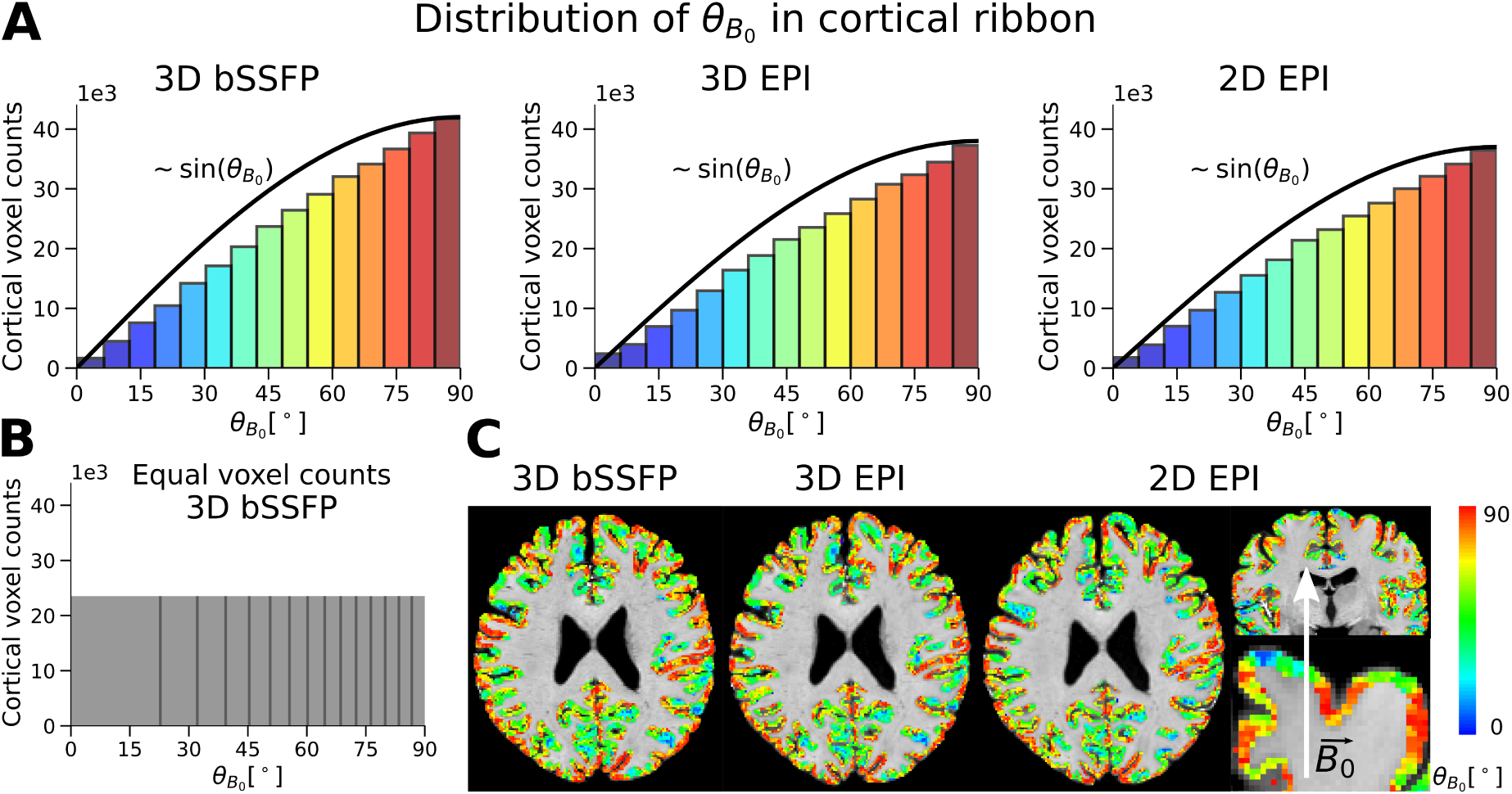
Distribution of θ_B0_ in the entire cortical ribbon after co-registration (and warping) (A). It can be seen that the distribution follows a sine curve. In order to perform the analysis without a higher weighting of voxels around 90°, the orientation intervals are adjusted as shown in (B) for further analysis. The co-registration (and warping) results for the orientation values for all three sequences are visualized (C). In addition to the axial slice (through-plane 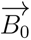 vector) a coronal slice and a zoomed view for the 2D EPI sequence is shown with the 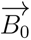 vector indicated as a white arrow.

### 3.2 Cortical Orientation and Cortical Depth Dependence

The cortical depths, *θ_B_*_0_, and the CV values in the functional space were used to compare the signal fluctuation of all three sequences with respect to their dependence on both the cortical orientation to 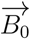 and the cortical depths. In order to have matching spatial coverage of 36 slices for all three sequences, the outer most slices were removed symmetrically (2, 4 and 25 from 3D bSSFP, 3D EPI and 2D EPI, respectively). For each cortical region of interest, the mean of the CV values of all runs within each (adjusted) cortical orientation bin, 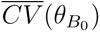, is normalized to the corresponding mean within the *θ_B_* ∼ 90*^◦^* bin, yielding the relative CV:

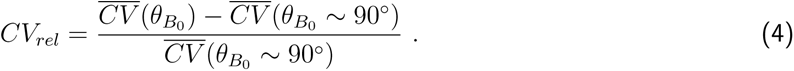

This normalization is performed as the lowest signal fluctuation is theoretically expected and found in voxels with *θ_B_* ∼ 90*◦* (vessels parallel to 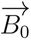).

Voxels from similar slice positions from all subjects were investigated and results from all sequences in session 1 are shown in Figure 5, which displays the relative CV vs. cortical orientation in the entire cortical ribbon (first row) and in each of the five cortical depths (rows 2 through 6). Different markers denote different subjects and the mean of all five subjects is shown in blue. Due to the adjusted *θ_B_*_0_ bin size per subject, each plotted point includes equal voxel numbers, as shown in Figure 4B. The mean *θ_B_*_0_ over all sequences plotted in the cortical ribbon to get equal voxel counts per bin were as follows: 14.04, 26.57, 34.41, 40.90, 46.68, 52.01, 57.00, 61.68, 66.09, 70.29, 74.27, 78.04, 81.61, 85.04, 88.36. Plotted CV values for the different sequences in the different cortical depths can be found in Supplementary Table S1.

**Figure 5:**
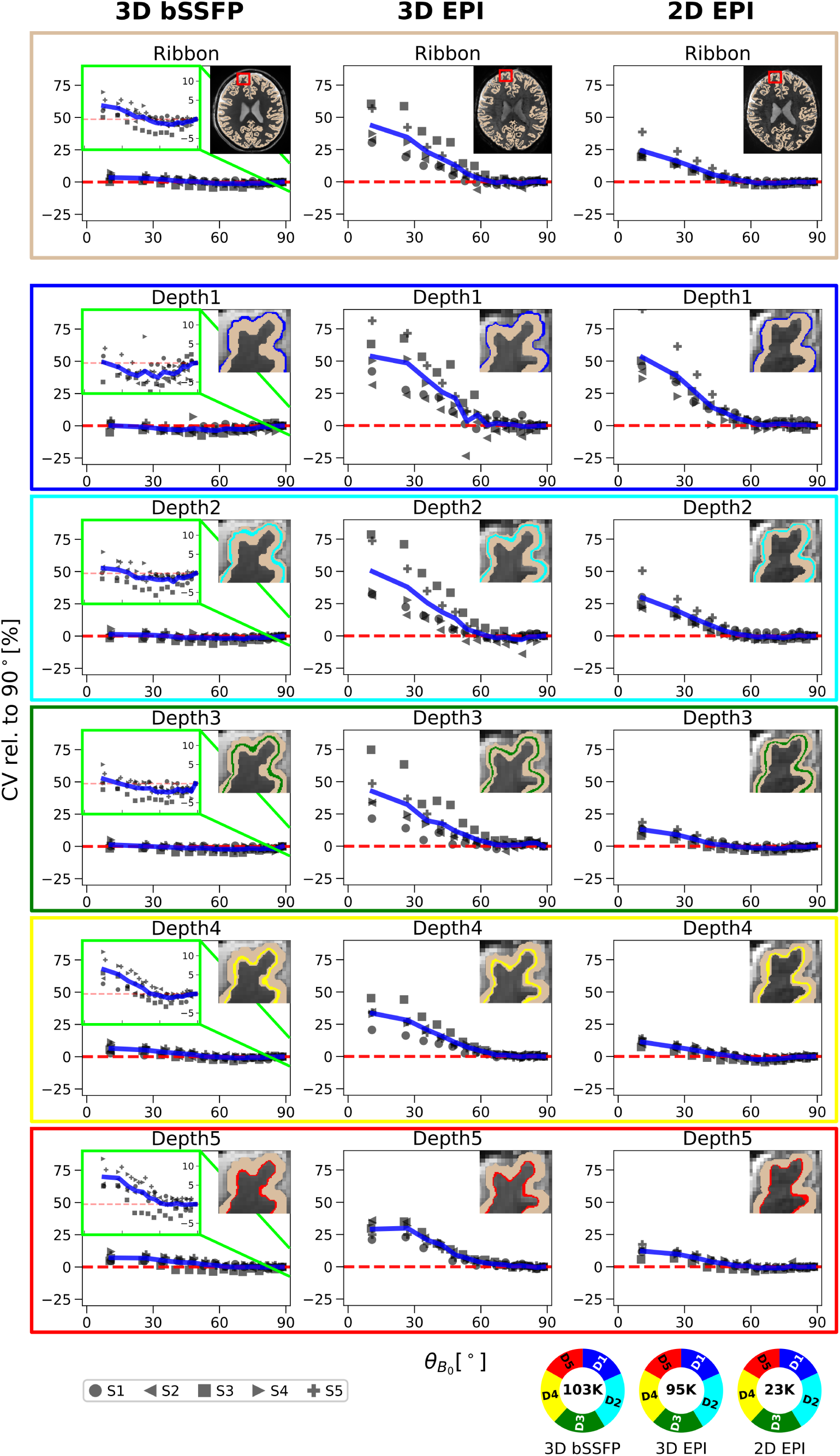
*Relative coefficient of variation in the up-sampled data in respect to the cortical orientation to* 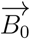 *for 3D bSSFP (left), 3D EPI (middle) and 2D EPI (right):* The mean CV_rel_ values calculated from all runs of each subject in session 1 are plotted against *θ_B_*_0_ in the entire cortical ribbon (top row) and in each cortical depth (rows 2 - 6). *θ_B_*_0_ is defined as the angle between the cortical surface normal and 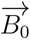. Depth 1 and Depth 5 are closest to CSF and WM, respectively. The cortical depth is also visualized in a zoomed-in exemplary slice of one subject in each plot. The CV values are plotted relative to the value at *θ_B_* = 90*^◦^* (CV_rel_)indicated by the red dashed line at 0%. The blue line shows the mean CV_rel_ across all subjects. Due to the lower variability of the CV in bSSFP, zoomed-in details are shown in the green boxes for a decreased range of 8 % to 14 %. The pie chart at the bottom shows the average number of data samples per point in the cortical ribbon plot across subjects. The distribution of voxels in each cortical depth is represented by the colors.

For both EPI sequences a strong dependence of the relative CV values on *θ_B_*_0_ is visible. Furthermore, the largest relative CV values and the strongest inter-subject variability especially in the shallow cortical depths, can be seen for 3D EPI. In the bSSFP data, no such dependence is observed and the CV values are barely affected by the cortical orientation at any depth. The inset plot for bSSFP (green frame) shows the relative CV values on a much smaller scale (in the range of −8 % to 14 %) across the entire orientation range to appreciate the small orientation dependence that is yet present.

In Depth 1 (closest to the cortical surface), the mean CV value over all subjects at approximately *θ_B_*_0_ = 14*^◦^* is 0.19 %, 53.59 %, and 53.06 % greater than the value at approximately *θ_B_*_0_ = 90*^◦^* for 3D bSSFP, 3D EPI and 2D EPI, respectively. At Depth 5 (closest to WM), this value increases slightly to 7.16 % for 3D bSSFP, whereas a decrease in orientation dependence can be seen for 3D EPI (29.16 %) and 2D EPI (12.12 %). Thus, the largest decrease in orientation dependence of CV for increasing cortical depths is noticeable in the 2D EPI plots (decrease of almost 41 %). The mean and standard deviation of the time series of all sequences plotted on *θ_B_*_0_ can be found in the Supplementary Figure S1.

The pie chart in Figure 5 shows the mean number of data samples across subjects for each plotted point in the entire cortical ribbon. For the analysis, only the data acquired during the first session are considered, which explains the lower number of voxels for each plotted point in the cortical ribbon for 2D EPI (one run) compared to the 3D sequences (four runs). The inverse number of runs was acquired in session 2, where a comparable behavior for the different cortical orientation dependence of CV can be seen (Supplementary Figure S2).

To further investigate the depth dependence observed in Figure 5, the mean of all CV values of each session is plotted against the cortical depths without normalization to *CV* (*θ_B0_* ∼ 90). In Figure 6A the depth dependence of each sequence is shown as absolute values and relative to D5, which is closest to WM and should therefore show the least influence from large pial vessels. Since D5 also shows the least variance across orientations compared to the other depths, it is used for normalization in the second row of Figure 6A. Markers are color-coded depending on their cortical orientation, with lighter orange indicating higher *θ_B_* values. A tendency of *θ_B_* ∼ 90 to have a lower dependence on the cortical depths can be seen in both EPI plots. The highest dependence of the signal fluctuation on the cortical depth can be seen in the 3D EPI plot. For the 2D EPI, the dependence is much lower but a strong variability across cortical orientations is still visible. As already visible in Figure 5, the bSSFP data show much lower CVs and a reversed trend in depth dependence. In addition, the influence of the cortical orientation on CV appears much lower for bSSFP.

**Figure 6:**
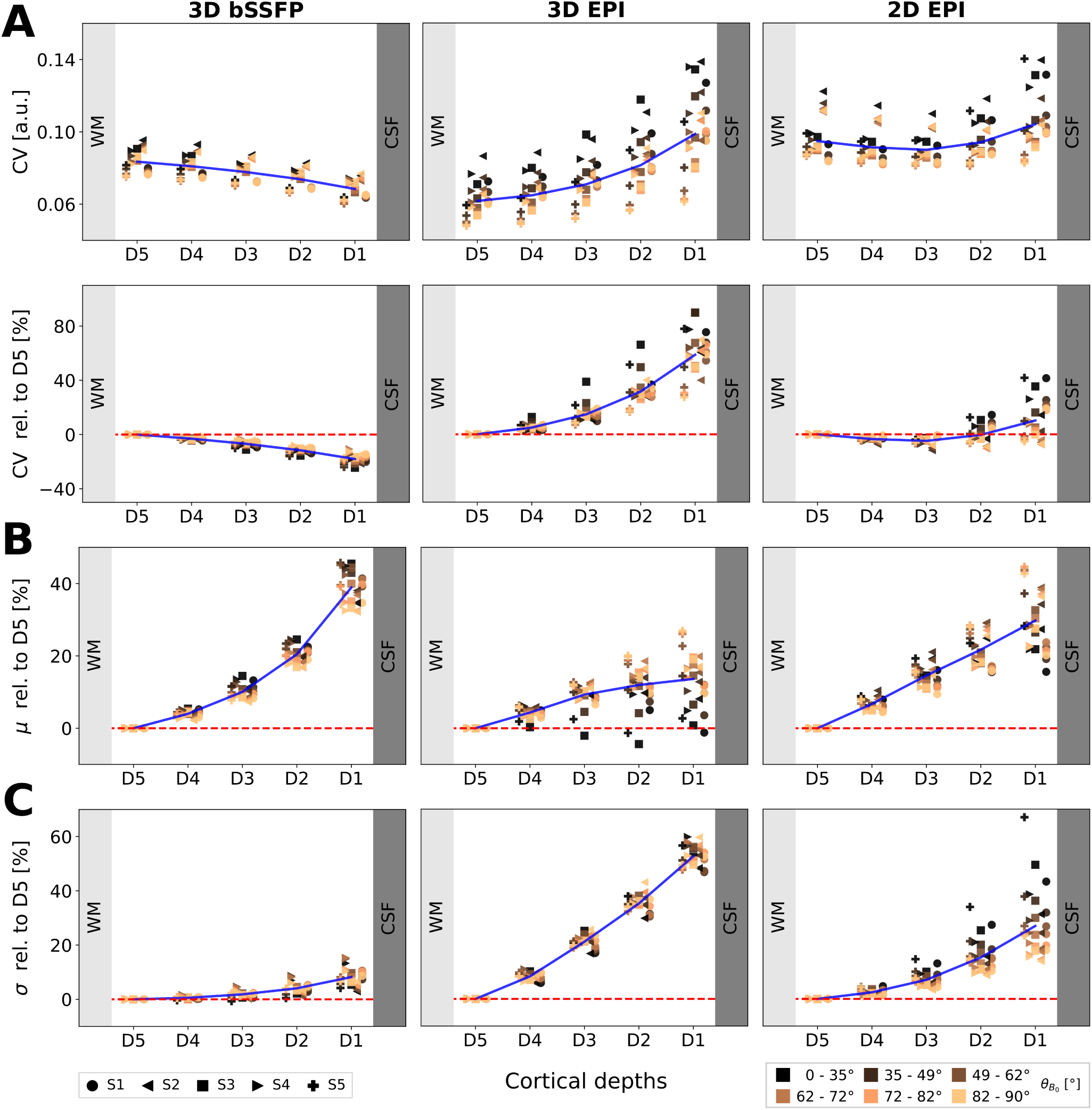
CV, µ and σ plotted against the cortical depth for 3D bSSFP (left), 3D EPI (middle) and 2D EPI (right). A: The absolute CV values (top) and relative to D5 (bottom) are plotted against the cortical depth with D1 and D5 closest to CSF and WM, respectively. Each point represents the mean of all CV values within a specific range of *θ_B_*_0_ values in one session. Six ranges are plotted for each subject, with lighter orange corresponding to higher *θ_B_*_0_ values around 90°, see legend. The intervals are not constant, so that each plotted point includes the same number of voxels for each subject. The mean of all subjects is shown in blue. The mean (B) and the standard deviation (C) of the time series are plotted similarly against the cortical depth.

The mean (signal amplitude) and standard deviation (signal variance) of the time series, that both contribute to CV, are plotted on the cortical depth (Figure 6B and C) relative to D5. For 3D bSSFP, the mean increases almost exponentially towards CSF and the standard deviation increases only slightly. For 3D EPI, the mean increases towards CSF in an orientation dependent manner, with *θ_B0_* ∼ 0 showing almost no increase. For one subject (S3), these values even decrease and then increase towards CSF. The average standard deviation is 52.6 % higher in D1 compared to D5. For 2D EPI, the mean signal increases almost linearly towards CSF without any dependence on the cortical orientation, while the standard deviation also increases towards CSF, but with a higher slope for *θ_B_*_0_ = 0*^◦^*than *θ_B_*_0_ = 90*^◦^* and a stronger inter-subject variability.

### 3.2 Cortical Orientation and Cortical Depth Dependence

### 3.3 Influence of Proximity to Large Veins

Co-registration and warping of the anatomical data and values calculated in the anatomical space to the functional data were most challenging for the ME-GRE data, as they were obtained during a different session. Co-registration results can be seen in Figure 7. The top image shows the calculated vein distance map with yellow closer and red further from a detected vein overlaid on the co-registered volume from the second echo. Below, the final co-registration (and warping) to all three sequences is shown. In GRE-EPI, veins appear dark, and it can be seen that the overlaid vein distance map follow the veins tightly in most locations.

**Figure 7:**
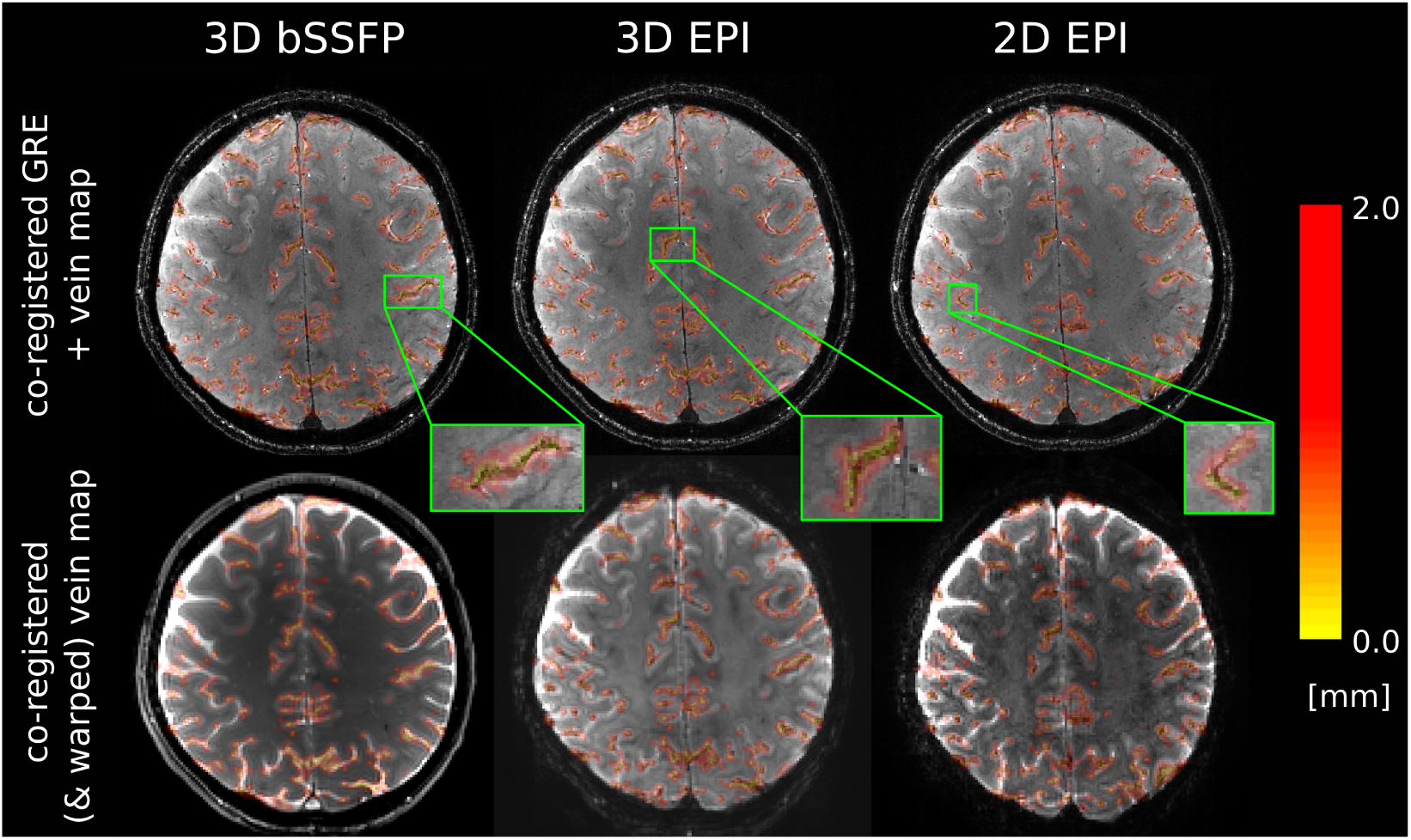
Vein distance map. shown on the co-registered gradient echo volume from which they were calculated (top), and overlaid on the mean functional images after co-registration (and warping) (bottom). Yellow voxels are closer to veins and red further away.

According to the shown results, it should be clear, that removing voxels in proximity to large draining veins will decrease the effect of cortical orientation dependence if a sequence is highly sensitive to these large veins. We show the cortical orientation dependence of three pools of voxels in Figure 8 for subjects S2-5 since the venous vessel mask was only available from these subjects. The blue line includes voxels within detected veins and with distances up to 0.71 mm (high proximity) from the veins. The green and the red lines include voxels, at 0.71 mm to 1.41 mm (medium proximity) and 1.41 mm to 2.12 mm (low proximity) distance from veins, respectively.

**Figure 8:**
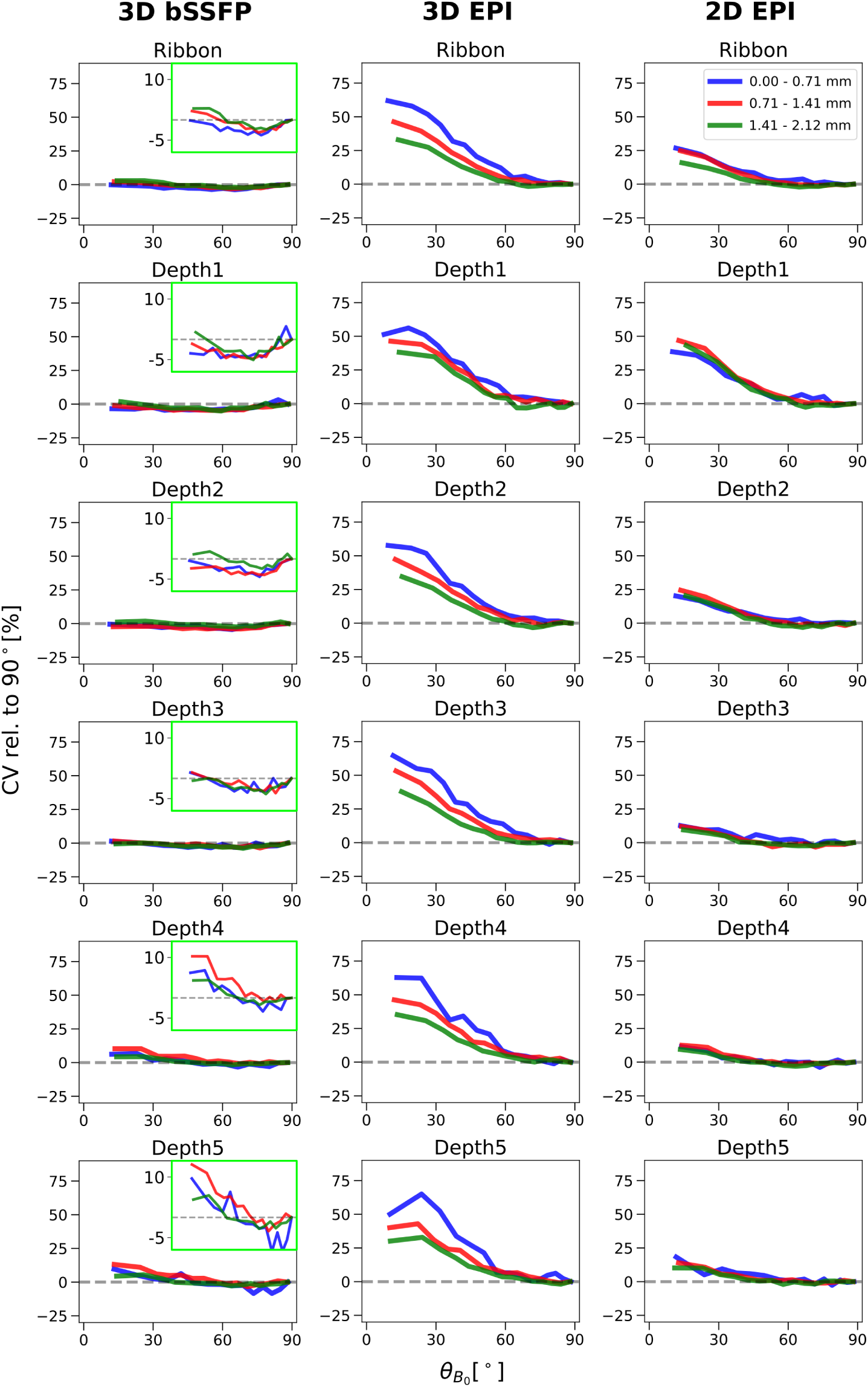
*CV dependence on the cortical orientation to* 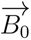 *in three distances to veins.* Distances to segmented veins in the range of 0 mm to 0.71 mm, 0.71 mm to 1.41 mm and 1.41 mm to 2.12 mm are represented by the blue, red and green lines, respectively. The mean CV_rel_ of four subjects is plotted in equally sized bins on *θ_B_*_0_ in the entire cortical ribbon and in each cortical depth. Zoomed-in versions of the bSSFP plots are shown in green boxes, similar to Figure 5.

The effect of vein distance on the cortical orientation dependence is particularly remarkable in 3D EPI. When looking at the pool of voxels further away from the veins, the cortical orientation dependence decreases. This can not only be seen in the entire cortical ribbon, but also in each cortical depth. In the cortical ribbon, the CV value at approximately *θ_B_*_0_ = 14*^◦^* is 61.89 % higher than the value at *θ_B_*_0_ = 90*^◦^* for voxels in high proximity to veins. This value is reduced by 15.67 % and 28.98 % for voxels in medium (red) and far distance (green) to the veins, respectively. Only when looking closely, a decrease of orientation dependence is also visible in 2D EPI, and this only in the entire cortical ribbon (when averaging all depths). Here, the value at *θ_B_*_0_ = 14*^◦^* closest to veins is 26.74 % higher than the *θ_B_*_0_ = 90*^◦^*. This value is decreased by only 1.95 % and 10.85 % for voxels at medium and low proximity, respectively. In bSSFP, this effect is slightly reversed, with the lowest value shown for voxels closest to veins. The value at approximately *θ_B_*_0_ = 14*^◦^* is 0.15 % lower than the value at *θ_B_*_0_ = 90*^◦^* for voxels closest to veins. This is increased by 2.34 % and 2.99 % for voxels in medium and low proximity, respectively.

In addition, absolute CV values and CV values normalized to D5 from voxels binned in the same manner as explained above were plotted in respect to the cortical depth (Figure 9). As expected, the CV values exhibit a greater increase towards the cortical surface when voxels are in closest proximity to veins for both EPI sequences. In 3D EPI, the highest signal fluctuation is seen for voxels of D1 when closest to veins. This effect is reduced, but is still prominent for voxels with distances 1.41 mm to 2.12 mm from the apparent vessel surface. In 2D EPI, the green curve (furthest from veins) even almost becomes flat. This clearly demonstrates that the high signal fluctuation in the EPI sequences on the cortical surface is partly of venous origin. Still the slope in 3D EPI is significantly higher than in 2D EPI when moving away from the veins. In contrast, the 3D bSSFP plot across cortical depths does not change for any of the voxel pools. Whether voxels are close to a vein or further away, the pattern of CV dependence is almost identical and varies little across cortical depths, with the lowest CV found in voxels located on the cortical surface.

**Figure 9:**
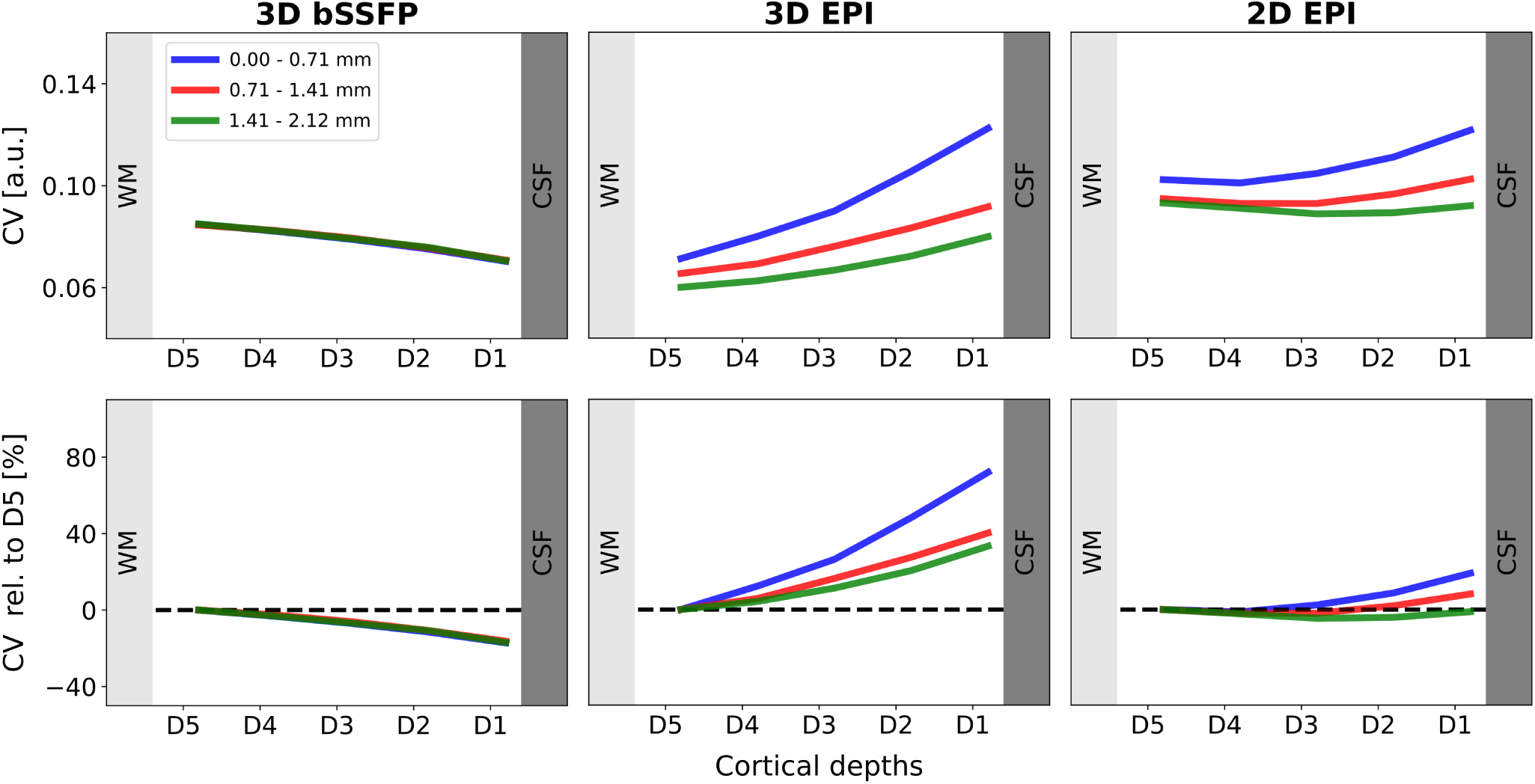
CV dependence on the cortical depth in three distances to veins. Vein distances of 0 mm to 0.71 mm, 0.71 mm to 1.41 mm and 1.41 mm to 2.12 mm are represented by the blue, red and green lines, respectively. The mean CV of four subjects is plotted against the cortical depth. For more details see caption of Figure 6A.

In Figure 10 we show the CV of all three sequences plotted as a function of the vein distance (absolute and relative to the value at smallest vein distance). It is expected, that voxels closer to the veins fluctuate more, thus have a higher CV value for the T2*-weighted sequences. For 2D and 3D EPI, the absolute CV values are highest when closest to large veins, which is not the case for 3D bSSFP. This once again highlights the sensitivity of the GRE-EPI to large veins, which is not seen for bSSFP.

**Figure 10:**
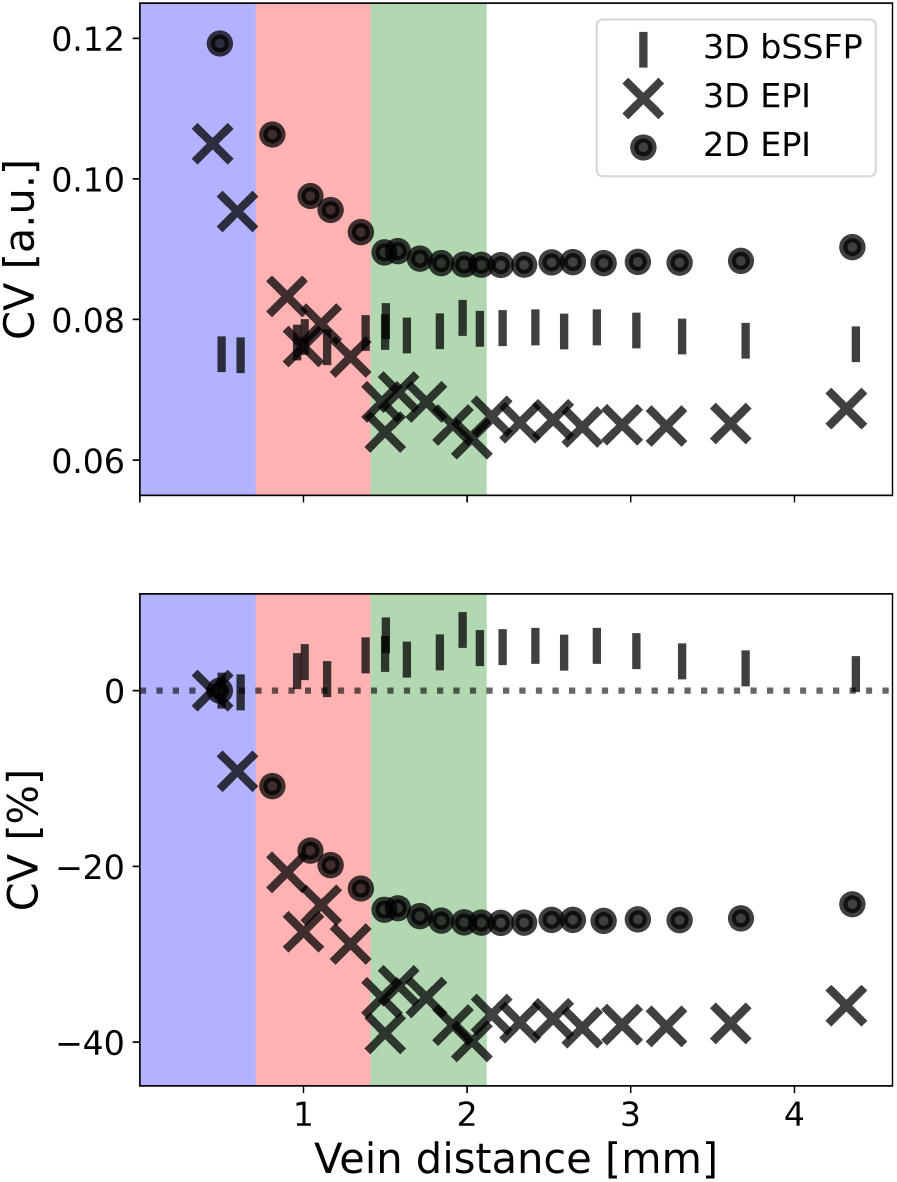
CV plotted on vein distance for 3D bSSFP, 3D EPI and 2D EPI. The absolute values (top) and relative to the voxels at highest proximity to veins (bottom) are shown. Vein distance = 0 mm are voxels inside visible veins in the SWI reconstruction. Data points contributing to Figure 8 and Figure 9 are highlighted in the same color used there. Within a sequence, every marker represents the same number of voxels (340 K, 320 K and 300 K for 3D bSSFP, 3D EPI and 2D EPI, respectively).

### 3.4 Inter-session Stability

Until here, we only show the cortical orientation (Figure 5, Figure 8) and cortical depth (Figure 6, Figure 9) dependence of the signal fluctuations in one session. The results from the second session are shown in Supplementary Figure S2 and Figure S3 (with vein distance in Supplementary Figure S4 and Figure S5). To investigate the repeatability of our observations, we plot the mean CV values of one run from session 2 against the mean CV values of one run from session 1 for all subjects (Figure 11). To this end, the coefficient of variation for thirty ranges of *θ_B_*_0_ (indicated by color) were extracted from one run of each session and plotted against each other. Each cortical angle range has the same number of voxels. The EPI plots show a high repeatability between the different sessions. This is also expressed in the correlation coefficients for individual subjects (Table 1), where a high repeatability between the sessions can be seen, especially for 2D and 3D EPI. However, the bSSFP plots show a lower repeatability, which might be explained by the small range of CV values (shown as a gray box in all plots). Thus, the correlation coefficients of the CV as function of *θ_B_*_0_ for bSFFP only ranges between 0.77 and 0.96 compared to 0.94 to 0.98 for the two EPI sequences. Nevertheless, the obtained correlation coefficients values indicate a high repeatability of the cortical angle dependence of the CV within subjects and sequences.

**Figure 11:**
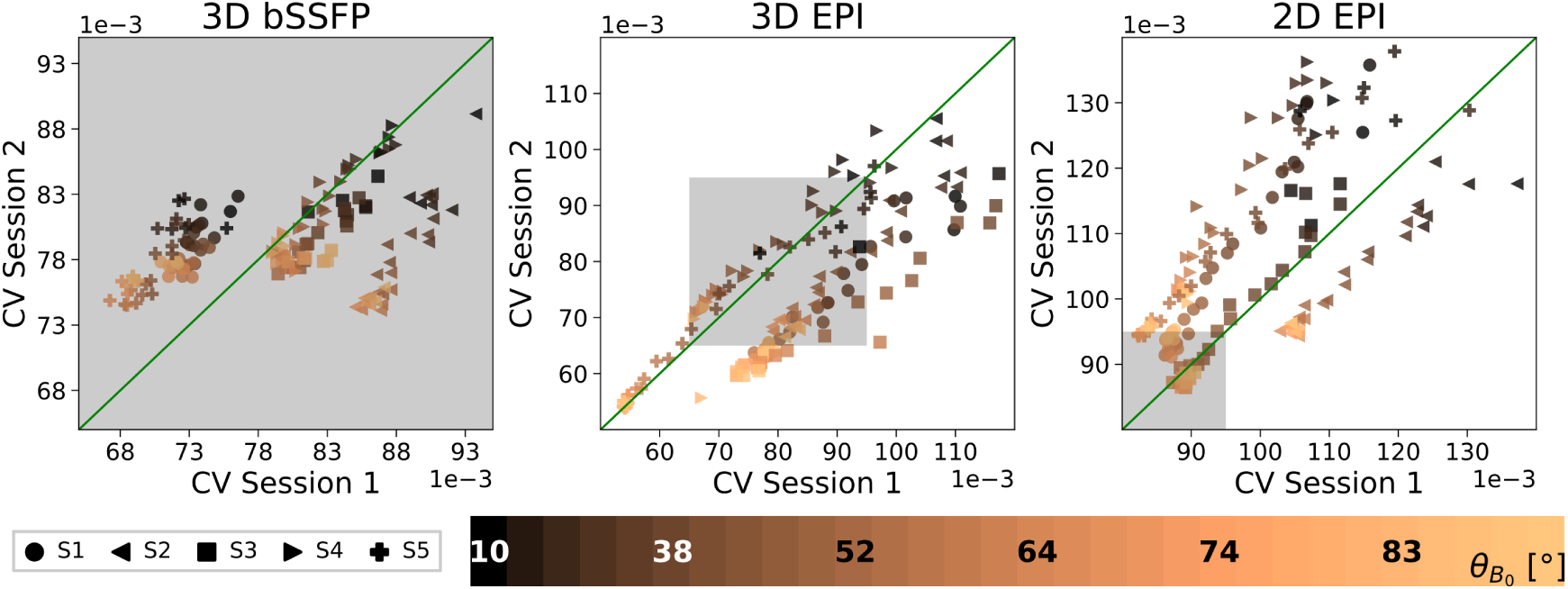
Inter-session variability. Mean CV values from run one of session 2 are plotted against mean CV values from run one of session 1 for all subjects. Thirty *θ_B_*_0_ ranges are shown for each subject with their mean *θ_B_*_0_ value shown in the color bar for every fifth range. The green line represents the ideal case, where both sessions are indistinguishable. Correlation coefficients are listed in Table 1. Note that each of the sequence plots has different x and y ranges. The range shown in the bSSFP plot is highlighted in the EPI plots as a gray box.

**Table 1:**
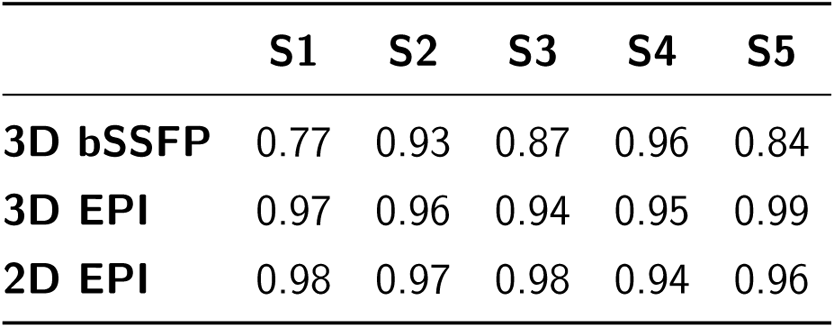
*Correlation coefficient* of the CV values plotted in Figure 11 calculated for each subject and sequence between sessions 1 and 2 using the first run of each.

|  | S1 | S2 | S3 | S4 | S5 |
| --- | --- | --- | --- | --- | --- |
| **3D bSSFP** | 0.77 | 0.93 | 0.87 | 0.96 | 0.84 |
| **3D EPI** | 0.97 | 0.96 | 0.94 | 0.95 | 0.99 |
| **2D EPI** | 0.98 | 0.97 | 0.98 | 0.94 | 0.96 |

## 4 Discussion

In this study, coefficient of variation values of resting-state fMRI signals from three sequences acquired across multiple runs and subjects were compared at 9.4 T. Specifically, the dependence on a) the cortical orientation relative to 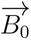, b) the cortical depth and c) their distance to veins was investigated. To this end, we first examined these influences on 2D SMS EPI at 9.4 T with the same parameters used by Viessmann *et al*. at 7 T (Viessmann et al., 2019) to compare our results with the existing literature. Imaging parameters of the 3D EPI and 3D bSSFP sequences like coverage, temporal and spatial resolution as well as excitation pulse duration and bandwidth-time-product were set to nearly identical values in order to be able to minimize potential bias. Although results from both EPI sequences were not congruent, a strong dependence of their signal fluctuations on cortical orientation was observed, allowing for a relative comparison between EPI and bSSFP signal fluctuations. According to our results, the cortical orientation to 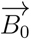 has almost no effect on the signal fluctuations from the bSSFP sequence. The observed CV variation is at least one order of magnitude smaller and follows different curves as a function of cortical orientation, cortical depth and vein distance compared to the EPI sequences. This may be an indicator of a higher spatial specificity of the bSSFP signal, showing that the signal is sensitive to the microvasculature and not to large pial veins on the cortical surface responsible for this effect. The following sections will discuss factors that might affect the results and comparison to existing literature.

### 4.1 Methodological Challenges and Considerations

Here, we investigate the rs-fMRI signal of three sequences. The global brain activation in rs-fMRI is advantageous for this work’s objective, as it allows larger parts of the brain to be studied simultaneously compared to typical task-fMRI paradigms, limited only by sequence and acquisition parameters. Another way to obtain large scale changes of blood oxygenation signal changes could be through hypercapnic tasks such as breath-holding or even light breathing challenges using moderate levels of O_2_ reduction / CO_2_ increase (Bright et al., 2009; Kastrup et al., 1998; Pinto et al., 2021). However, due to subject specific breathing patterns, as well as increased subject movement during deep inhalation and exhalation required for the breath hold task, such experiments are usually difficult to perform and evaluate. Furthermore, hypercapnia might not only lead to a change of oxygenation in the vascular level, but also complex physiological changes like decreased metabolism and neuronal activity which makes the interpretation of signal changes even more complex (Peng et al., 2017). We tried to reduce influences on the rs-fMRI signal by measuring four runs of five minutes each, giving us 20 minutes per subject for each sequence. But even if we compare two sessions taken at the same time of day, with different numbers of runs (session 1: 4 x 3D sequences & 1 x 2D sequence; session 2: 1 x 3D sequences & 4 x 2D sequence), we get similar results. Nevertheless, it would be interesting to investigate in the future, how the BOLD signal e.g. from a visual task is affected by cortical orientation to 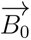 using the same sequences used in this study. The challenges would include the smaller brain region being investigated and – in the example of the visual cortex – the limited distribution of cortical orientations in that region.

To make both 3D sequences comparable, the TR_vol_ and the voxel size were kept constant, while adjusting the acceleration factors for parallel imaging. Since a higher acceleration factor of R = 5 was used for the 3D bSSFP sequence compared to the segmented 3D EPI with R = 3, higher g factor losses can be expected for this sequence which would increase noise variance between voxels in resulting images (Avdievich et al., 2019). However, CV values reflecting the reciprocal of tSNR were comparable between the sequences (see Figure 2 and absolute CV values in Figure 6), enabling a valid comparison of their dependence on the cortical orientation and cortical depth. Another point of discussion regarding image acquisition could be the effect of variations in the excitation flip angle due to transmit field inhomogeneities, a common problem in UHF MRI. In this study, all measurements except the MPRAGE scans were performed with a static phase shift between the transmit coil elements corresponding to a circularly polarised mode. Thus, for all rs-fMRI scans, the deviations from the nominal flip angle are comparable within subjects and across sessions. We therefore believe that the impact of this effect on the presented results is very limited. Like Viessmann *et al*. (Viessmann et al., 2019) we chose the temporal CV as a measure of signal fluctuation. To be able to compare absolute CV values of all three sequences to each other, we additionally normalized it by the 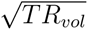, as the 2D sequence was acquired with a TR of 1.7 s (see Viessmann et al., 2019) and the 3D sequences were acquired with a TR of 3 s. For the major part of our results, however, any global scaling factors were eliminated by using relative CV metrics (with respect to CV at *θ_B_*_0_ ∼ 90*^◦^*, with respect to CV at D5, or with respect to CV at the smallest vein distance).

Other factors that may have influenced our results include the segmentation of the brain into GM and WM masks calculated with FreeSurfer, which were then used to divide the cortical band into five equally spaced depths. The co-registration of brain masks and cortical orientation values to the functional space also poses a critical step in the analysis. Volumes co-registered to the EPI space had an additional step of warping to match the EPI distortions. More precisely, while the (readout) distortions of MPRAGE and bSSFP were negligible, such that they matched well natively, the (different and non-negligible phase encode) distortions of both EPI data required additional warping of the cortical band before layerification. This step introduced additional interpolation bias compared to the distortion free bSSFP that cannot be accounted for easily. Therefore, after warping the anatomical tissue masks, the data were visually inspected to ensure a good fit to the functional images.

The cortical orientation values (*θ_B_*_0_) were calculated from the T1-weighted MPRAGE, which was acquired in the same session as the shown results for all subjects except subject S1. If *θ_B_*_0_ were calculated from a different session, this might add a bias for rotations around the x and y axes. Due to the dimensions of the used coil (Shajan et al., 2014), excessive rotation around these axes was not possible. Nevertheless, the x and y rotations were calculated from the transformation matrices during co-registration. As expected, maximum rotation occurred, when the MPRAGE image was acquired in a different session and was at Δ*θ_B_* = 4.7*^◦^*for subject S5 between session 1 and 2. Since the visualisation was done by averaging over irregular ranges of *θ_B_*_0_ to keep the number of data points in each range constant, a small deviation like Δ*θ_B_* = 4.7*^◦^*is hardly noticeable. We furthermore show the results of session 2 of all subjects in Supplementary Figure S2 and Figure S3, which still pointed to the same effects, even when the *θ_B_*_0_ values were calculated in a different session. The intersession stability in Figure 11 and the correlation coefficients in Table 1 were further indicators, that calculating *θ_B_*_0_ in different sessions did not affect the outcome.

### 4.2 2D EPI Cortical Orientation and Cortical Depth Dependence

Viessmann *et al*. investigated the signal fluctuation dependence of 2D SMS EPI on the cortical orientation at 7 T (Viessmann et al., 2019). An even higher effect is expected at 9.4 T, as the susceptibility effect and thus the draining vein bias of a GRE-EPI sequence becomes higher with increasing field strengths. Although similar parameters were used to acquire the 2D EPI data, a direct comparison with the existing literature is not straightforward. This is due to the fact that Viessmann *et al*. used FreeSurfer for depth segmentation from superficial CSF through GM to subcortical WM. Furthermore, the authors performed temporal smoothing, and presented the data without removing the orientation bias from equal orientation bin sizes (Figure 4A). We used LayNii for depth segmentation of GM only, did not perform temporal smoothing, and present the data excluding the orientation bias (Figure 4B). Although the inclusion of CSF and WM voxels theoretically allows for the investigation of the CV in regions with different vascular architecture and where no or little functional signal is expected, such additional layers are difficult to define unambiguously due to partial volume effects. Temporal smoothing might be beneficial to reduce thermal noise. However, we abstained from modifying the temporal signal to not introduce any potential interpolation bias to the data. Still, the effect of temporal smoothing and physiological noise reduction was investigated for one subject and can be seen in Supplementary Figure S6 and Figure S7. Regressing out physiological noise was not feasible for all runs of all subjects, due to an error during data collection, but has been done for one example subject (see below). Finally, we present our data without the orientation bias, by adjusting the sizes of the orientation bins (Figure 4B), facilitating their interpretation without unequal CV precision across cortical orientations.

Nevertheless, despite the different analysis approach, the cortical orientation dependence of 2D EPI persists at 9.4 T and was comparable to the effects observed at 7 T. Highest CV values occurred for *θ_B0_* → 0*^◦^* (pial vessels perpendicular to 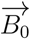) and decreased towards *θ_B_* = 90*^◦^* (Figure 5). Due to the removal of the orientation bias in our data, fewer data points were plotted around *θ_B_* ∼ 0_0_. Therefore, it cannot be determined, whether our data follow a cos^2^(*θ_B_*) curve down to *θ_B_* = 0, as expected and shown by Viessmann *et al*.. However, it was also not completely ruled out by our data. While the orientation dependence was present in all cortical depths, it decreased closer to WM. This is expected for 2D EPI (and 3D EPI), as gradient echo EPI is T2*-weighted and highly biased by the extravascular signals caused by large pial veins on the cortical surface. The CV of 2D EPI decreased slightly and then increased towards CSF (Figure 6A), as similarly shown by Viessmann et al., 2019. It was also shown that CV values from regions with lower *θ_B_*_0_, corresponding to greater perpendicularity of the pial veins, show a greater slope across the cortical ribbon. This is expected when veins are perpendicular to 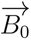, due to the maximization of their signal contribution. In addition, previous theoretical and experimental studies have shown that large pial veins are responsible for signal bias even at greater cortical depths (Bause et al., 2020; Koopmans et al., 2010; Pais-Roldàn et al., 2023; Polimeni et al., 2010; Ress et al., 2007; Siero et al., 2011).

### 4.3 3D EPI vs. 2D EPI

Compared to 2D EPI, a higher cortical orientation dependence was observed for 3D EPI signal fluctuation (Figure 5). Both sequences however showed a CV decrease towards WM, reflecting the reduced influence of the large veins near WM compared to the cortical surface. When comparing signal fluctuations from both sequences at different cortical depths (Figure 6), 3D EPI showed a higher increase towards the cortical surface, than 2D EPI, except when the cortex was perpendicular to 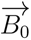.

One explanation for the observed discrepancies between 2D and 3D EPI might be that 3D EPI is known to be more susceptible to physiological influences (heart rate and breathing) than (single-shot) 2D EPI (Lutti et al., 2013) due to repeated excitation of the entire slab for Fourier encoding along the slice direction. While 3D bSSFP also employs repeated slab excitations and Fourier slice encoding, 3D EPI combines this with strong T_2_*-/susceptibility-weighting (long TE), which leads to image artifacts that show as time series signal fluctuations. This might also explain the high variance in the 3D EPI data (Figure 6C). After regressing out a major source of physiological image noise for one exemplary subject using RETROICOR (Glover et al., 2000), the orientation dependence is reduced both for 2D and 3D EPI, but the CV dependence remains larger for 3D EPI (Supplementary Figure S6, Figure S7). Another physiological noise correction approach (RVHR), that is more directly linked to vascular oxygenation changes via respiratory variations (RV) and variations of the heart rate (HR), may be interesting to investigate in this context (Chang et al., 2009). However it is known that the respiratory and cardiac response functions employed in such corrections vary across brain regions. A careful analysis, that is beyond the scope of the work, would therefore be needed to avoid yet another potential cortical orientation bias. Furthermore, we observed stronger signal fluctuations and dynamic geometric distortions in the inferior section (closer to the neck) of the 3D EPI scans which is indicative of its physiological origin. In comparison, movements of tissue borders were not visible in the 2D EPI measurements and the mean of the time series shows an almost linear increase towards the cortical surface (Figure 6B). This indicates that another typical BOLD time series correction step not performed in this work, i.e. regression of motion parameters (and their derivatives) to remove signal variations explainable by motion (Power et al., 2014), may change the overall fluctuation amplitude differently in 2D and 3D EPI. There is also a difference in the readout bandwidth, coverage and parallel imaging between the two sequence since the 2D SMS-EPI protocol was designed such that a comparison to the data of Viessmann et al., 2019 can be performed. In contrast, the 3D protocol was optimized with regard to coverage, temporal resolution and tSNR of the bSSFP sequence.

Other parameters that have a potential effect on the CV include TE and echo train duration (*τ*). In our case, however, we kept both of these parameters almost constant between the EPI sequences. Moreover, images were acquired with different TR_vol_ in both sequences (1.7 s and 3 s for 2D and 3D EPI, respectively). To explore the impact of this discrepancy, an additional analysis was performed by dropping every second volume in 2D EPI, resulting in an effective TR of 3.4 s. After a comparison between the resulting plots, no difference was seen (results not shown). This rules out the possibility that TR_vol_ might be responsible for the +19.54% increase of CV of 3D EPI in the entire cortical ribbon compared to 2D EPI.

In summary, physiological influences or different motion sensitivities might explain the observed differences between the amplitudes of 2D and 3D EPI cortical orientation and depth dependence. Further work is, however, needed to carefully analyze these EPI differences without introducing orientation bias, whereas this work focuses on the differences to the 3D bSSFP sequence.

### 4.4 3D bSSFP Cortical Orientation and Cortical Depth Dependence

While a clear and high dependence of CV and thus the signal fluctuation of the EPI signals on the cortical orientation relative to 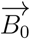 was observed, this was not the case for bSSFP, especially in the shallow depths. Surprisingly, the signal fluctuation was similarly dependent on *θ_B_*_0_ in the deeper cortical depths (D4 and D5), but with reduced magnitude compared to both EPI sequences (Figure 5).

The extravascular effect of veins at the GM-WM boundary, which often drain deep cortical layers and run parallel to the cortex before ascending to the cortical surface (Duvernoy et al., 1981) may explain the cos^2^ (*θ_B_*) dependence seen in Depth 4 and Depth 5. Since the signal dependence on orientation cannot be described by a sin^2^ (*θ_B_*) at any cortical depth, it can be assumed that the bSSFP measurements were not sensitive to ascending veins, or only to a very limited extent.

If both, surface veins and orthogonal intracortical veins contribute equally to the signal, no dependence on the cortical orientation to 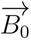 is expected. The same is the case if bSSFP rs-fMRI signals are predominantly sourcing from randomly oriented capillaries where, in addition, almost 50 % of the signal is intravascular and 50 % extravascular (Pérez-Rodas et al., 2021). This would negate any dependence on the vein orientation. We observed this behavior in the zoomed-in version of the bSSFP plots for D2 and D3 (green boxes in Figure 5).

In accordance with the aforementioned observations, the cortical depth dependence plots (Figure 6) showed a decrease of the coefficient of variation towards the cortical surface for bSSFP in contrast to both EPI sequences. On the contrary, Bàez-Yàñez *et al*. show that the BOLD signal increases towards the cortical surface for all simulated sequences, regardless of whether it is GRE, SE, or bSSFP. Although this is not consistent with our findings regarding the relationship between the coefficient of variation (CV) and cortical depths, we observe the same effects when plotting the standard deviation of the time series against the cortical depths. This means that the course of the CV relative to the cortical depths can only be explained by the much stronger increase of bSSFP signal towards the cortical surface.

The reduced cortical orientation dependence in bSSFP indicates that the signal contribution in bSSFP is not dominated by the extravascular component of the macrovasculature. This work would be the first experimental demonstration of the bSSFP’s sensitivity to the microvasculature, which had previously been hypothesized based on Monte Carlo simulations (Bàez-Yànez et al., 2017; Pérez-Rodas et al., 2021; Scheffler et al., 2019). A reduced macrovascular signal bias would make bSSFP a good alternative for measuring the BOLD signal at high and UHF, e.g. in order to investigate cortical depth dependent signals in laminar studies.

### 4.5 Comparison with Simulation Studies

Bàez-Yàñez *et al*. simulated the vessel orientation effect on GRE, SE and bSSFP at 9.4 T (Bàez-Yànez et al., 2017). They showed the highest dependence on the vessel orientation for GRE, especially for large draining veins (Figure 4 in Bàez-Yànez et al., 2017). In the SE simulations a generally lower dependence was shown, with the highest dependence experienced by the microvasculature (≤10 µm). They furthermore show, that this effect in bSSFP was highly dependent on the chosen TR and FA. From the simulated parameters, TR / FA = 5 ms / 20*^◦^* were the closest to our chosen parameters (TR / FA = 3.14 ms / 11*^◦^*). In this case, their bSSFP results showed a similar effect as experienced by the SE sequence; an overall reduced dependence on the cortical orientation. Additionally, signal change from the microvasculature (≈1 µm to 10 µm) showed a higher dependence on the vessel orientation than vessels with diameters of 50 µm to 200 µm, which was reversed for the GRE sequence. Unfortunately, the effect of the randomly oriented microvasculature cannot be investigated with our method, as they would cancel each other out. However, we were able to show the lack of orientation dependence in the depths closest to CSF, highlighting the negligible signal contribution from the large draining veins on the cortical surface in bSSFP.

Nevertheless, the above-mentioned signal simulations of Bàez-Yàñez *et al*. did not exactly match our results (see their Figure 8) which may be due to several reasons: First, the BOLD effect was simulated, meaning that two states were compared, while we were looking at the fluctuations of the resting-state signal. Second, the vascular network model used in their simulations was based on vessel segmentation of the mouse parietal cortex. It is however known that the ratio of arteries to veins is reversed between mice and humans and that the human cortical vascular network varies much more across brain regions. This is why efforts are being made to generate human vascular artificial networks to simulate the BOLD signal more realistically (Cassot et al., 2006; Hartung et al., 2022; Linninger et al., 2019; Lorthois et al., 2011). Third, in the simulations, deep cortical depths were simulated purely as such, without the effect of partial volumes and potential inflow from WM regions. Fourth, because the WM/GM boundary is not identifiable in the vascular models, it is not clear whether the models used in the simulations cover the entire cortex or only parts of it. Finally, the included pial veins in the mouse vascular model are much smaller with diameters up to 70 µm (according to Figure 5 in Bàez-Yànez et al., 2017) compared to ranges of 50 µm to 200 µm in humans (Duvernoy et al., 1983). Thus, in humans they can be even much smaller than the resolution used for the generation of the vessel mask in this work which further complicates the comparison of our calculated vessel distance effects to the simulated ones. For these reasons, it is difficult to compare signal simulation using non-human vascular models with our results.

### 4.6 Influence of Large Veins

Our results suggest that moving away from large veins detected in the SWI images reduces the orientation dependence experienced by the signal fluctuation in GRE-EPI. If the effect of the large surface veins were completely eliminated, the dependence on the cortical orientation would have to be reversed (∼ sin^2^ (*θ_B_*)) due to the dominance of signals sourcing from orthogonal ascending veins. However, this effect is not seen in our plots, which means that merely removing voxels inside veins or close to veins will not result in an elimination of their effect on the signal (fluctuation), especially at the used spatial resolution. Previous work also showed that the effects of the large veins (≥ 0.3 mm in diameter) can induce perturbations, which are perceptible even 6.7 mm away from the vein (Huck et al., 2023). Much smaller veins are probably not detectable with our method (Bause et al., 2020; Bollmann et al., 2022). Although the dependence on the cortical orientation relative to 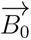 was not reversed when moving away from the veins, it was diminished. These findings further enhance the potential of our method to be used for studying the macrovascular contribution to the fMRI signal.

### 4.7 Layer fMRI at UHF

Care should be taken when performing resting-state layer fMRI with GRE-EPI sequences, as it is extremely biased by the large draining veins (Bause et al., 2020; Turner, 2002). This is not only the case at UHF, and efforts exist to remove the large veins contribution, even at 3 T (X. Z. Zhong et al., 2024). In this work, we were able to reproduce the observations on the rs-fMRI signal originating from the large draining veins through analysis of the cortical orientation relative to 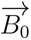 in all cortical depths. Compared to 2D and 3D EPI, 3D bSSFP signal fluctuations were only slightly affected by the cortical orientation and in a reversed way. They also did not show any dependence on the distance to the veins, reflecting its potential sensitivity to the microvasculature. Vascular space occupancy (VASO) (Lu et al., 2003) has been used as an alternative to the conventional BOLD imaging at UHF (Huber et al., 2014). Although direct measurements of cerebral blood volume changes using VASO or cerebral blood flow measurements can provide a much higher signal specificity than GRE-BOLD, they usually suffer from a low contrast-to-noise ratio, increased SAR and lower temporal resolution limiting their applicability for certain research questions. Based on our results, we believe that the bSSFP sequence may be another good alternative to conventional GRE-EPI sequences, allowing depth-dependent functional and resting-state fMRI without signal bias due to cortical orientation and geometric distortions. However, future work is needed to assess the signal specificity of bSSFP and EPI sequences using task-based layer fMRI and to improve BOLD signal modelling, e.g. by using realistic artificial vascular networks.

## 5 Conclusion

In this study, we have investigated the influence of a) cortical orientation, b) cortical depth, and c) vein distance on resting-state fMRI signals of three different sequences: 3D bSSFP, segmented 3D GRE-EPI, and 2D SMS GRE-EPI. While both GRE-EPI sequences exhibited a high cortical orientation dependence, an elevated signal change towards the cortical surface, and a reduced orientation and depth dependence with increased distance to veins, this was not the case for 3D bSSFP. Thus, all three examined parameters (a-c) indicate reduced macrovascular contributions in rs-fMRI scans with bSSFP compared to GRE-EPI and therefore a higher specificity for signal changes originating from small veins within the cortex, closer to neuronal activity. Our work provides the first experimental confirmation of the signal characteristics of bSSFP, previously estimated from Monte Carlo simulations. This makes bSSFP a promising sequence for layer fMRI applications, especially at UHF.

## Data and Code Availability

Anonymized and defaced MRI data used in this project will be made available upon reasonable request.

## Author Contribution

The authors confirm contribution to the paper as follows: study conception and design: JB, DR; supervision of research: KS; data collection: DR, JB, SM; analysis of data: DR; technical support: SM, RS, DB, PE; interpretation of results: DR, JB, KS; draft manuscript preparation: DR. All authors reviewed the results, edited the manuscript and approved its final version.

## Acknowledgements

We thank our coil engineers for keeping the coil running.

