## Supplementary Material for "Macrovascular contributions to resting-state fMRI signals: A comparison between EPI and bSSFP at 9.4 Tesla"

### 6 Supplementary Material

Table S1: Relative CV values plotted in Figure 5 using 15 bins with the same amount of voxels in each bin. Corresponding mean $\theta_{B_0}$ values are not shown here, but can be seen in the plots. R: entire cortical ribbon; D1-5: different cortical depths.

**3D bSSFP**

| | | | | | | | | | | | | | | | |
|---|---|---|---|---|---|---|---|---|---|---|---|---|---|---|---|
| R | 3.53 | 2.88 | 1.71 | 0.57 | 0.61 | -0.39 | -0.97 | -1.08 | -1.29 | -1.38 | -0.82 | -1.06 | -0.62 | -0.54 | 0.00 |
| D1 | 0.19 | -1.16 | -2.72 | -3.36 | -2.11 | -3.22 | -3.79 | -2.32 | -2.63 | -2.86 | -2.23 | -0.84 | -1.08 | -0.20 | 0.00 |
| D2 | 1.35 | 0.93 | 0.20 | -1.22 | -1.18 | -1.31 | -1.68 | -1.75 | -1.46 | -2.25 | -1.05 | -1.30 | -0.88 | -0.62 | 0.00 |
| D3 | 1.28 | 0.08 | -0.53 | -1.09 | -1.27 | -1.25 | -2.15 | -2.11 | -2.28 | -2.08 | -1.44 | -2.06 | -1.55 | -2.06 | 0.00 |
| D4 | 6.43 | 5.12 | 3.37 | 2.43 | 1.51 | 0.10 | -0.16 | -0.57 | -0.97 | -1.02 | -0.59 | -0.74 | -0.32 | -0.10 | 0.00 |
| D5 | 7.16 | 6.86 | 4.73 | 3.52 | 3.24 | 1.70 | 1.26 | 0.23 | -0.04 | -0.07 | 0.21 | -0.10 | -0.12 | 0.00 | 0.00 |

**3D EPI**

| | | | | | | | | | | | | | | | |
|---|---|---|---|---|---|---|---|---|---|---|---|---|---|---|---|
| R | 43.52 | 34.50 | 23.21 | 17.66 | 12.66 | 6.21 | 2.85 | 0.94 | -0.77 | 0.45 | -0.44 | -1.39 | -0.07 | 0.35 | 0.00 |
| D1 | 53.59 | 48.36 | 35.49 | 25.88 | 21.06 | 3.35 | 8.35 | 0.33 | 1.75 | 0.96 | 0.81 | -0.57 | -0.85 | -0.30 | 0.00 |
| D2 | 49.97 | 37.83 | 25.70 | 18.30 | 13.54 | 5.10 | 2.48 | 0.01 | -1.52 | -1.14 | -2.49 | -3.02 | -0.70 | -0.54 | 0.00 |
| D3 | 42.52 | 32.19 | 19.82 | 17.45 | 11.70 | 8.24 | 4.89 | 2.89 | 1.03 | 0.73 | 1.01 | 1.19 | 2.43 | 2.42 | 0.00 |
| D4 | 33.49 | 28.12 | 20.64 | 15.65 | 10.83 | 7.36 | 5.01 | 3.13 | 1.15 | 1.19 | 0.45 | -0.14 | 0.45 | 0.34 | 0.00 |
| D5 | 29.16 | 29.95 | 20.85 | 16.75 | 9.27 | 6.64 | 4.09 | 2.71 | 2.04 | 1.51 | 0.46 | 0.87 | 0.34 | -0.56 | 0.00 |

**2D EPI**

| | | | | | | | | | | | | | | | |
|---|---|---|---|---|---|---|---|---|---|---|---|---|---|---|---|
| R | 23.98 | 15.88 | 10.48 | 5.71 | 3.62 | 1.39 | 0.39 | -1.05 | -0.87 | -1.09 | -0.67 | -0.28 | -0.02 | -0.31 | 0.00 |
| D1 | 53.06 | 38.17 | 24.72 | 14.03 | 9.88 | 5.54 | 3.70 | 0.81 | 0.41 | 1.19 | 1.38 | 0.63 | 0.74 | 0.72 | 0.00 |
| D2 | 29.57 | 19.35 | 12.49 | 7.70 | 3.78 | 1.39 | 0.35 | -0.98 | -0.41 | -0.95 | -1.53 | -0.44 | 0.56 | -0.44 | 0.00 |
| D3 | 12.81 | 8.79 | 4.83 | 1.63 | 0.80 | -0.55 | -1.01 | -1.52 | -1.27 | -2.04 | -1.66 | -0.40 | -0.52 | -0.72 | 0.00 |
| D4 | 11.17 | 7.02 | 4.60 | 2.24 | 1.13 | -0.27 | -1.42 | -2.31 | -2.08 | -1.94 | -1.50 | -0.65 | -0.27 | -0.29 | 0.00 |
| D5 | 12.12 | 9.80 | 6.68 | 3.70 | 2.50 | 0.67 | 0.73 | -0.83 | -0.81 | -1.06 | -0.92 | 0.59 | -0.33 | -0.08 | 0.00 |

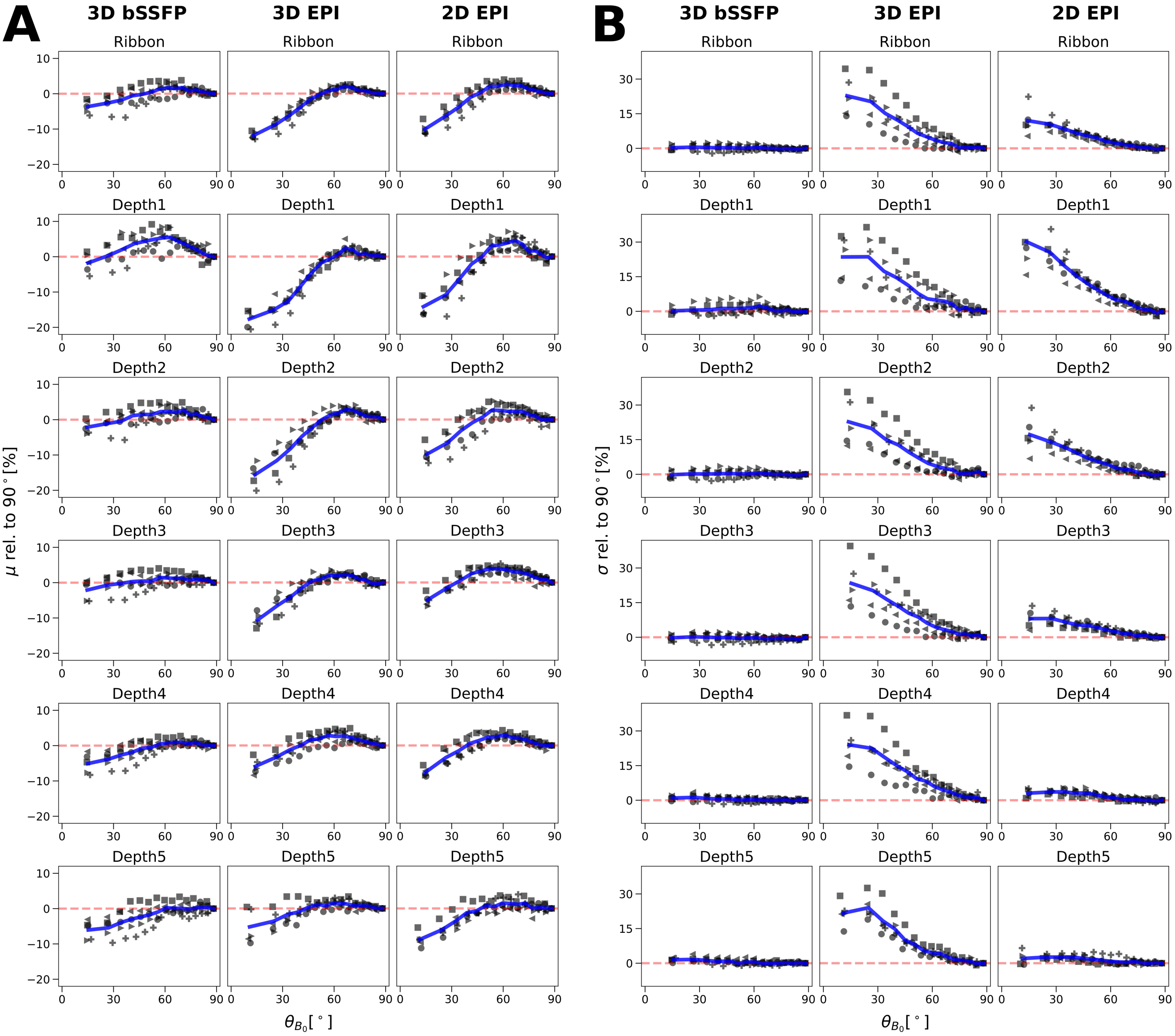


Figure S1: Mean **(A)** and standard deviation **(B)** of the time series plotted on the cortical orientation relative to B0. For details, see Figure 5.

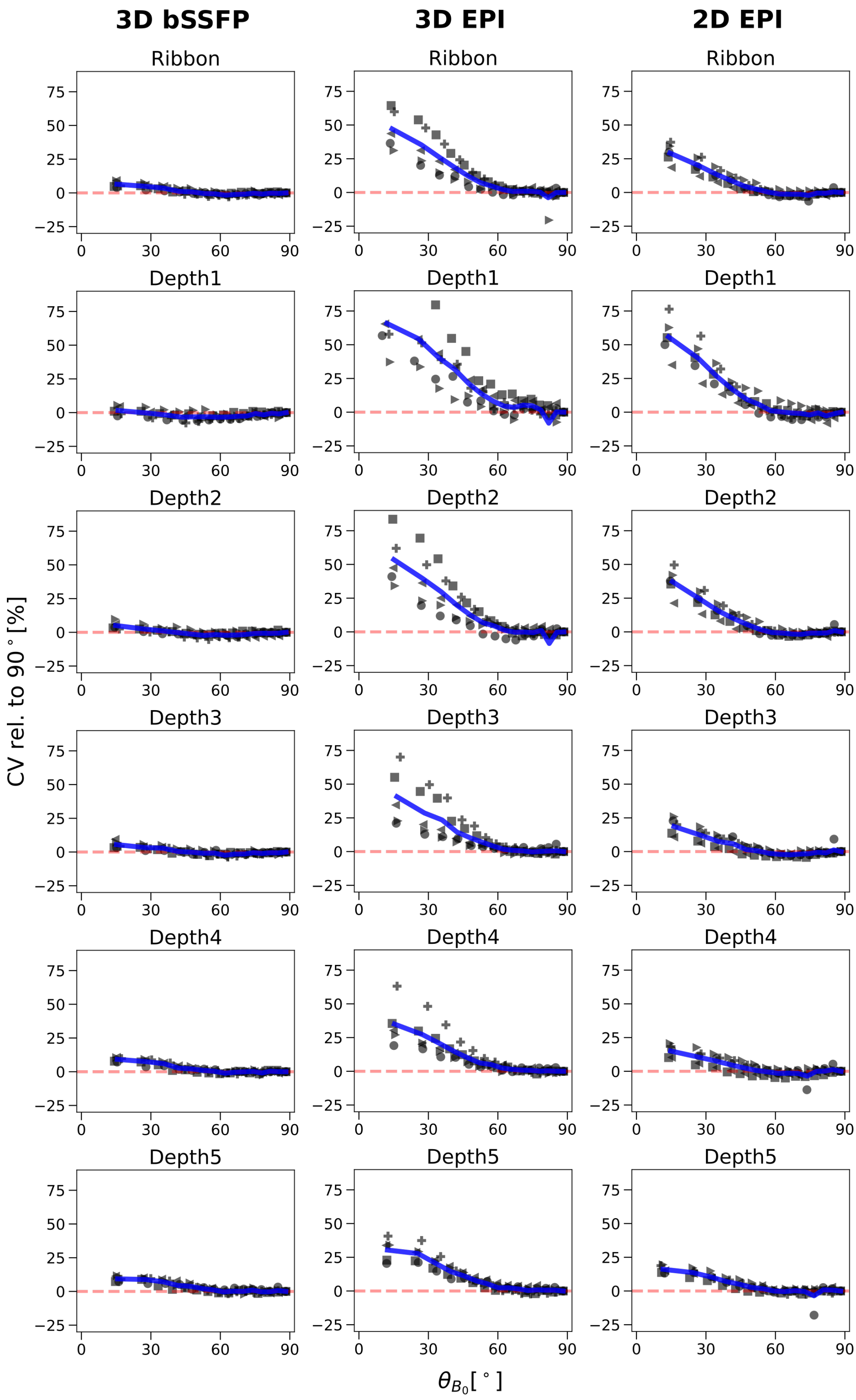


Figure S2: *$CV_{rel}$ plotted on the cortical orientation to B0 in session 2.* Mean $CV_{rel}$ values from one run (3D bSSFP and 3D EPI) and four runs (2D EPI) of all five subjects are shown in blue. See Figure 5 for details.

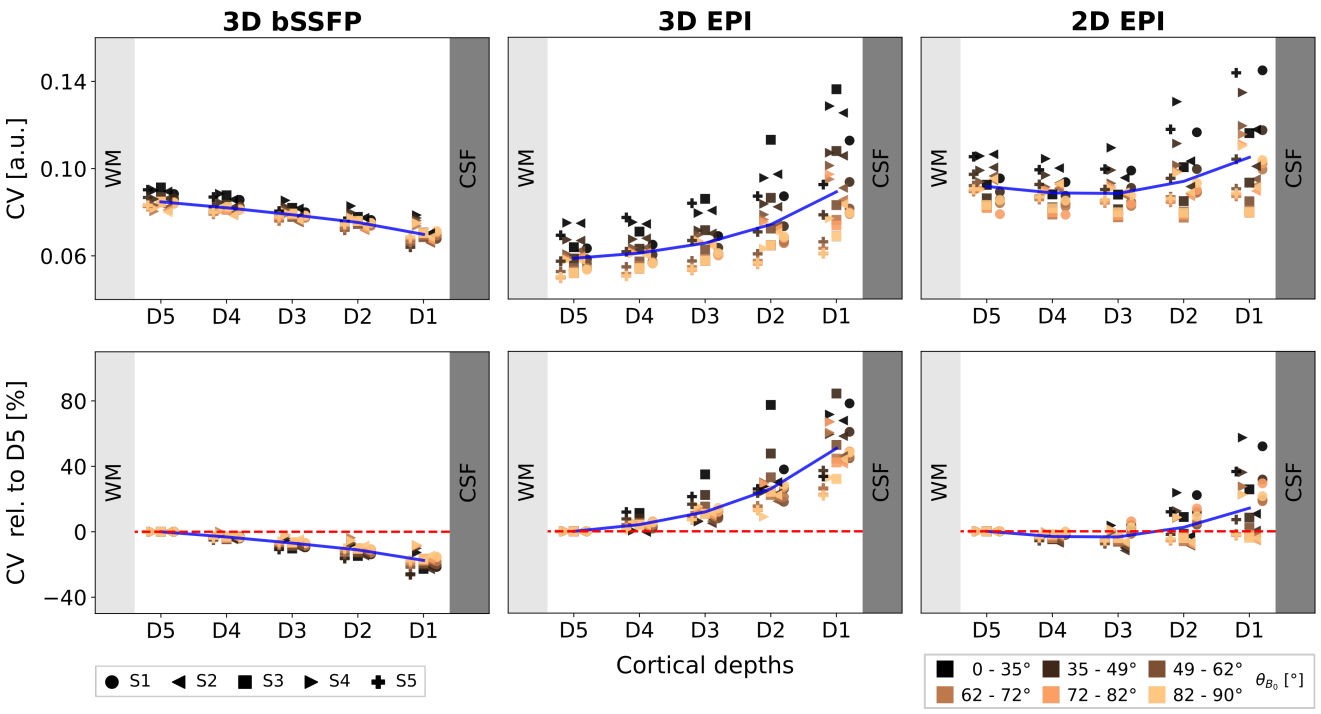


Figure S3: *CV plotted on the cortical depths in session 2* (D1 and D5 closest to CSF and WM, respectively). Mean $CV_{rel}$ values from one run (3D bSSFP and 3D EPI) and four runs (2D EPI) of all five subjects are shown in blue. See Figure 6 for details.

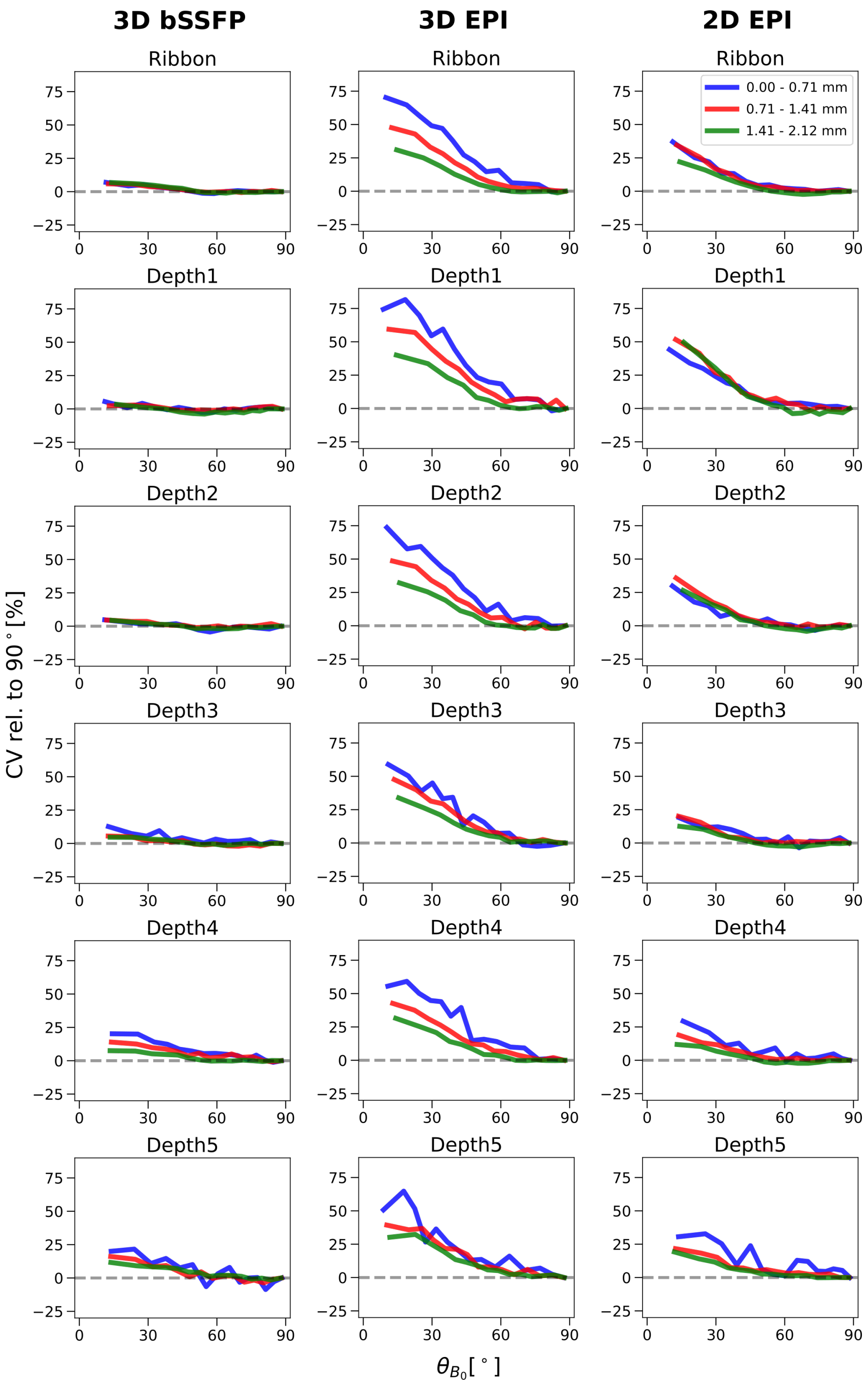


Figure S4: *$CV_{rel}$ plotted on the cortical orientation to B0 in session 2.* Mean $CV_{rel}$ values from one run (3D bSSFP and 3D EPI) and four runs (2D EPI) of four subjects are shown. Voxels are binarized into three pools depending on their distance from veins. The blue, red and green lines correspond to voxels in high ($0\,\mathrm{mm}$ to $0.71\,\mathrm{mm}$), medium ($0.71\,\mathrm{mm}$ to $1.41\,\mathrm{mm}$) and low ($1.41\,\mathrm{mm}$ to $2.12\,\mathrm{mm}$) proximity to the veins. See Figure 8 for details.

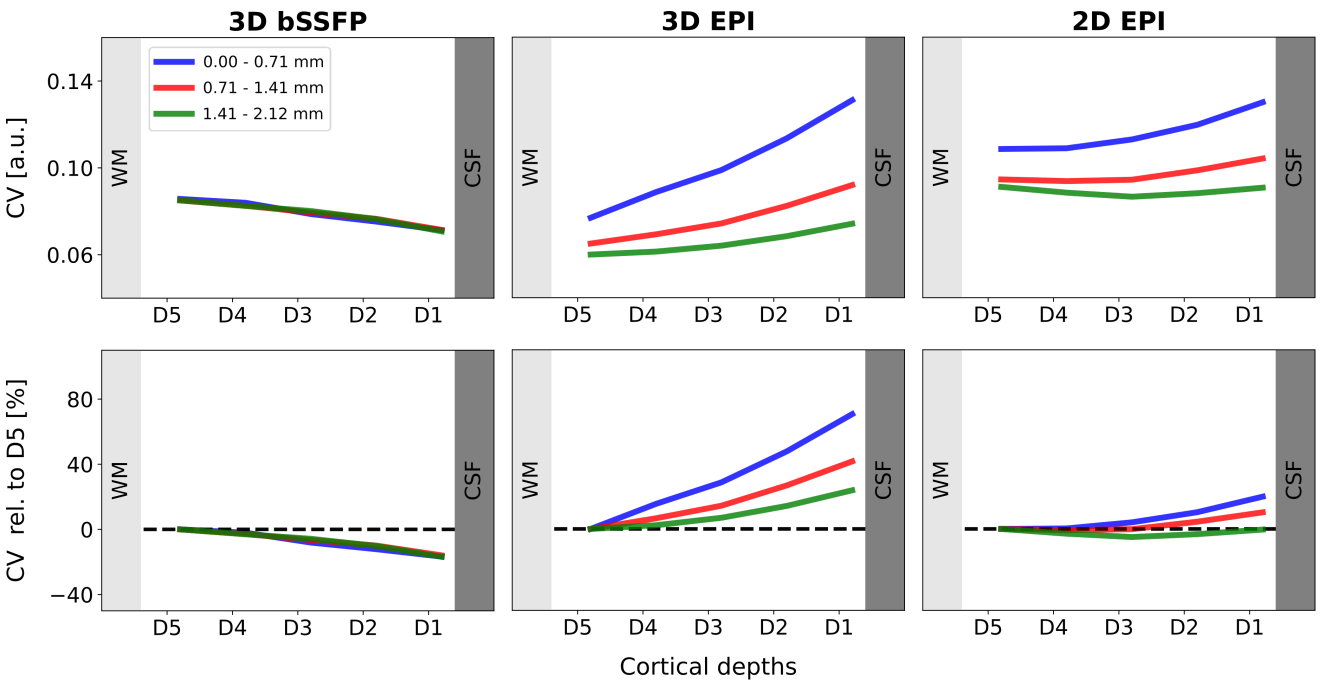


Figure S5: *CV plotted on the cortical depths in session 2.* Mean CV values from one run (3D bSSFP and 3D EPI) and four runs (2D EPI) of four subjects are shown. Voxels are binarized into three pools depending on their distance to veins. The blue, red and green lines correspond to voxels in high ($0\,\mathrm{mm}$ to $0.71\,\mathrm{mm}$), medium ($0.71\,\mathrm{mm}$ to $1.41\,\mathrm{mm}$) and low ($1.41\,\mathrm{mm}$ to $2.12\,\mathrm{mm}$) proximity to the veins. See Figure 9 for details.

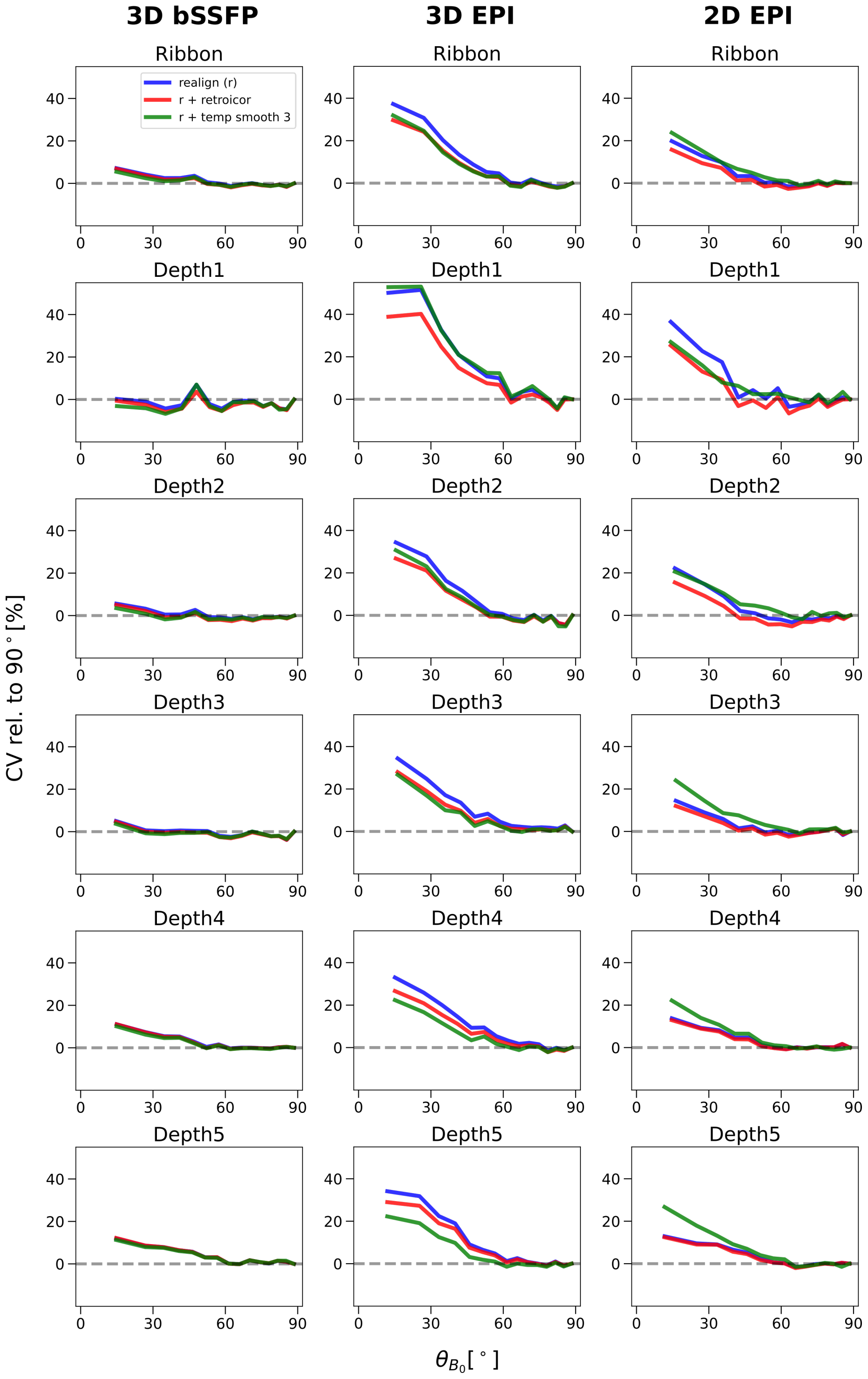


Figure S6: *Comparison of cortical orientation dependence after physiological noise regression and temporal smoothing:* $\mathrm{CV_{rel}}$ plotted $\theta_{B_0}$ for 3D bSSFP (left), 3D EPI (middle) and 2D EPI (right). The average $\mathrm{CV_{rel}}$ calculated after motion correction only [realign] **(blue)**, realignment and physiological noise regression using RETROICOR [r + retroicor] **(red)**, and realignment and temporal smoothing of the data with a moving average of 3 $\mathrm{TR_{vol}}$ [r + temp smooth 3] **(green)** are plotted on $\theta_{B_0}$. One subject (S4) is shown here, because the acquisition of the physiological parameters was only reliable for all runs of this subject. For more details, see caption of Figure 5.

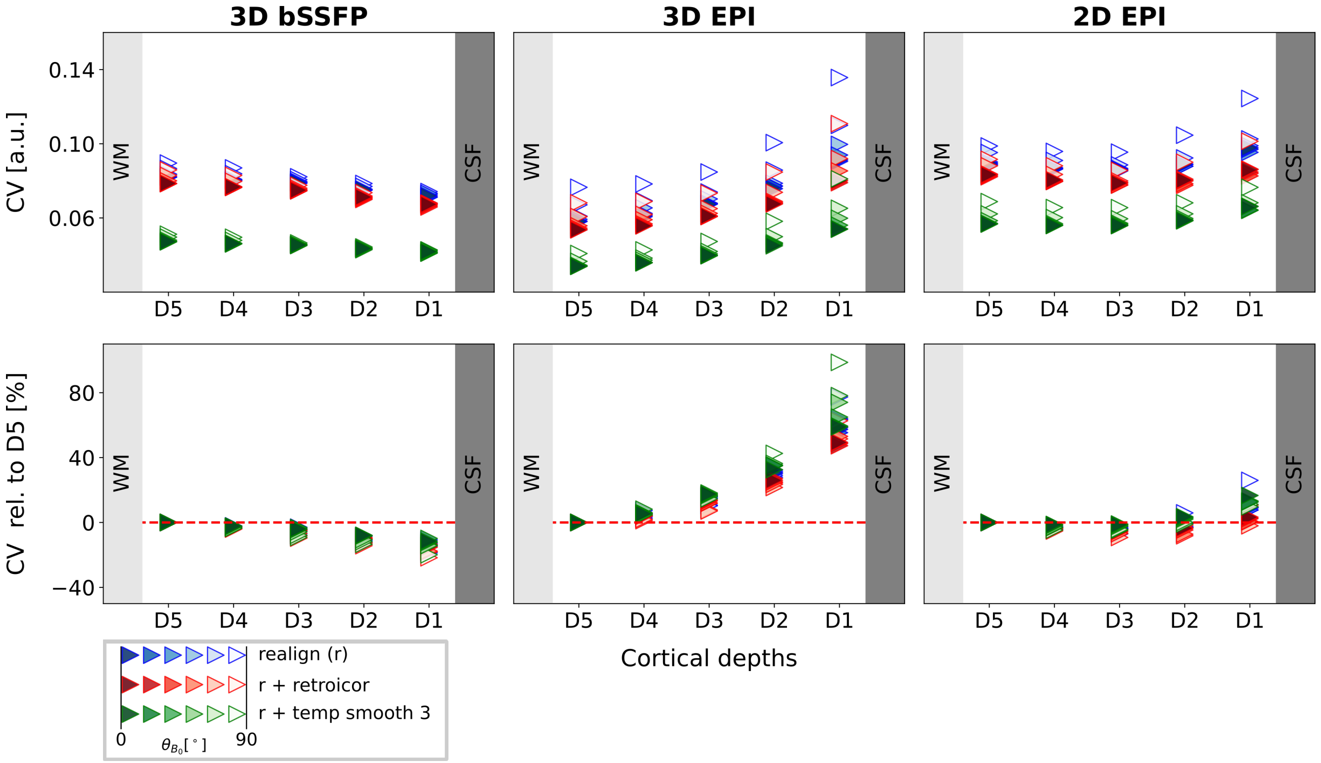


Figure S7: *Comparison of cortical orientation dependence after physiological noise correction using RETROICOR (red) and temporal smoothing with a moving average of 3 $TR_{vol}$:* The CV was plotted on the cortical depths for 3D bSSFP (left), 3D EPI (middle) and 2D EPI (right). The absolute (top) CV values and relative to D5 (bottom) are shown. Each plotted point represents the mean of all CV values within a specific range of $\theta_{B_0}$ values. Six ranges with equal voxel counts were plotted for each subject, with lighter colors corresponding to higher $\theta_{B_0}$ values around 90°. Only one subject (S4) is shown here, because the acquisition of the physiological parameters was only reliable for all runs of this subject.
